# Cell wall-resident PIR proteins show an inverted architecture in *Neurospora crassa*, but keep their role as wall stabilizers

**DOI:** 10.1101/2024.07.18.603779

**Authors:** Paul Montaño-Silva, Olga A. Callejas-Negrete, Alejandro Pereira-Santana, Jorge Verdín

**Author notes:** **Correspondence:** Jorge Verdín, Biotecnología Industrial, CIATEJ. Camino Arenero 1227, Zapopan, JAL, México 45019.

## Abstract

Proteins with internal repeats (PIRs) are the second most abundant class of fungal cell wall resident proteins. In yeasts, PIRs preserve the wall stability under stressful conditions. They are characterized by conserved N-terminal amino acid sequences repeated in tandem (PIR domains), and a Cys-rich C-terminal domain. Despite PIRs have been inferred in several filamentous fungi genomes, they have not been studied beyond yeasts. In this work, PIRs diversity, evolution and biological role, focused on a new PIRs class, were addressed. Bioinformatic inference of PIRs in fungi indicated they were an innovation in Ascomycota. Predicted PIRs clustered in two main groups: classical yeasts PIRs (N-terminal PIR domains; C-terminal Cys-rich domain), and PIRs from filamentous fungi with an inverted architecture (N-terminal Cys-rich domain; C-terminal PIR domains), which could harbor additional GPI-signals. As representatives of the second group, *Neurospora crassa* (Nc) PIR-1 (NCU04033) and PIR-2 (NCU07569) were studied. Confocal microscopy of eGFP-labeled PIR-1 and PIR-2 revealed they accumulate in apical plugs; additionally, PIR-1 requires the Kex2 processing site for correct maturation, and its predicted C-terminal GPI modification signal resulted functional. Moreover, Nc Δ*pir-1* and Δ*pir-2* single mutants showed a growth rate similar to that of Nc WT, but the double mutant Nc Δ*pir-1*/Δ*pir-2* grew significatively slower. Similarly, Nc Δ*pir-1* and Nc Δ*pir-2* were mildly sensitive to calcofluor white, although Nc Δ*pir-1*/Δ*pir-2* double mutant was severely impaired. Despite the inverted architecture of PIR-1 and PIR-2, they resulted in cell wall stabilizers as classical yeast PIRs.

## 1 INTRODUCTION

The cell wall is a sturdy but springy structure that surrounds the fungal cells protecting and adapting them to environmental stresses [1,2]. In general, the fungal cell wall is composed of chitin, chitosan, β-1,3-glucan, β-1,4-glucan, α-1,6-glucan, and cell wall resident proteins (CWP), either manno- or galactomannoproteins [2–4]. CWPs are the least studied components of the fungal cell wall. Several of them are enzymes that participate in the crosslinking and ripening of the wall [5–7], while others have been proposed as the cellular interface fungi need to get adapted to the environment [6,8]. Nevertheless, most of them remain uncharacterized and are ambiguously considered “structural” proteins [6].

After glycosylphosphatidylinositol-linked proteins (GPI-proteins), proteins with internal repeats (PIR) are the second most abundant covalently-bound CWPs [7]. In yeasts, PIRs are overexpressed under hypoxic conditions [9], confer heat-shock resistance and, in general, they are involved in the maintenance of the cell wall structure [10–13] by a β-1,3-glucan cross-connecting activity [7,14,15].

PIRs are defined by a series of N-terminal amino acid sequences repeated in tandem (PIR-domains) [16–18] that consist of eight highly conserved amino acids (SQGDGQQA) [19]. Variations to the PIR-domain have been suggested in *Saccharomyces cerevisiae* Scw4p and Scw10p glucanases [6,20]; however, their functionality remains to be determined. A single PIR may contain from one to up to 10 repetitive units, which defines the strength of the attachment to the wall [16–18,21]□. Additionally, they have a characteristic four cysteine domain at the C-terminus. PIRs can be covalently attached to the cell wall either through an alkali-labile ester bond between a glutamine residue of the repetitive units and β-1,3-glucans [19], or via a disulfide bond between the C-terminal Cys-rich domain and other CWPs [15,22]. Besides the arrangement of canonical PIRs, *S. cerevisiae* Cwp1p was found to contain both PIR-domains and a GPI-modification site, which are used simultaneously to bind it to the cell wall [23]. Because of their bimodal covalent attachment to the wall, PIRs have been often used as anchors for the display of enzymes on the yeast cell surface [21,24,25].

PIRs tandem repeats are part of a generalized characteristic among CWPs, which consistently contain repeated sequences in a wide range of number and length [26]. Protein repeats multiplication leads to a fast diversification of non-catalytic CWPs [8,27,28], which ultimately allows fungal cells to get adapted to a diversity of environmental stimuli.

PIRs have been widely studied in yeasts: *S. cerevisiae* [10,11,18], *Zigosaccharomyces rouxi* [12]□, *Kluyveromyces lactis* [29]*, Candida albicans* [30,31], *Yarrowia lipolytica* [17]□, *Pichia pastoris* [32] and *Candida glabrata* [33][. They have only been bioinformatically inferred for four filamentous fungi: *Magnaporte grisea, Gibberella zeae*, *Blumeria graminis* and *Neurospora crassa* [34]*. In N. crassa,* NCU04033 (PIR-1) and NCU07569 (PIR-2) loci encode putative PIRs. Unlike yeasts PIRs, *N. crassa* PIR-1 and PIR-2 show an atypical architecture; their PIR-domains are located close to the C-terminus, while only PIR-1 harbors a Cys-rich domain located close to the N-terminus [34]. To date, neither the diversity nor the function of PIRs in filamentous fungi have been studied. In this work, we aim to analyze the evolutionary history of PIRs in fungi and determine the functional role of these proteins in the cell wall using *N. crassa* PIR-1 and PIR-2 as models.

## 2 MATERIALS AND METHODS

### 2.1 Bioinformatic analysis

#### Mining of PIR-domain containing proteins

To identify PIR-domain containing proteins beyond Kingdom Fungi, Ensembl All [35], UniProtKB [36], and SwissProt [37] databases were thoroughly searched using the yeast PIR-domain (PF00399, downloaded from the Pfam 36.0 database [38]) as a query in the HMMER server [39] with an E<1×10^-4^. To assess the distribution of PIR-domain containing proteins in the Fungal Tree of Life, 840 fungal reference proteomes were downloaded (October, 2022) from the NCBI refseq [40] and scanned against the yeast PIR-domain (PF00399) in locally run HMMER, E<1×10^-4^. All retrieved sequences were manually curated, keeping only complete sequences. To avoid false positive PIRs, sequences were additionally filtered using a regular expression [QXXDGQXQ] that compelled the identification of amino acids necessary for cell wall anchoring within the PIR-domain [14].

Total putative PIRs were bioinformatically characterized: SignalP 6.0 [41] was used for signal peptide prediction; GPI signal identification was carried out with the NetGPI 1.1 predictor server [42]; transmembrane domains were predicted with deepTM located in the DTU/DeepTMHMM server [43]; PIRs serine/threonine composition was estimated with COPid [44]. To identify any additional functional domain, an hmmscan in HMMER server was performed for all PIRs (Pfam-A hmm database was used to search against it and 1×10^-4^ E-value was established as cut-off). IUPRED3 [45], FlDPnn [46] and ESpritz [47] were used for intrinsic disorder prediction.

For clustering analysis, EFI-EST server [48] was used according to the following parameters: fragment removal was set on, E-value was established as default (1×10^-5^), and minimum sequence similarity threshold was established as 46%. Reconstruction of the fungal tree of life was generated using 840 reference fungi species with PhyloT v2 (https://phylot.biobyte.de/), based on the NCBI taxonomy [49] and visualized and annotated using iTol v6 [128]. PIRs tertiary structure models were predicted using AlphaFoldv2, and oligomeric structures were predicted with AlphaFold3 server, both predictions using default parameters [50]. Figures were edited with the open source Inkscape v1.3.2 software.

### 2.2 Strains, media, culture conditions, and transformation

Bacterial and fungal strains used in this work are listed in Table 1. Bacteria were cultivated in liquid or solid Luria-Bertani medium at 37 °C [51] and supplemented with 1 µg/mL ampicillin when necessary. Ca-competent *E. coli* cells were used for heat-shock transformation.

**Table 1.**
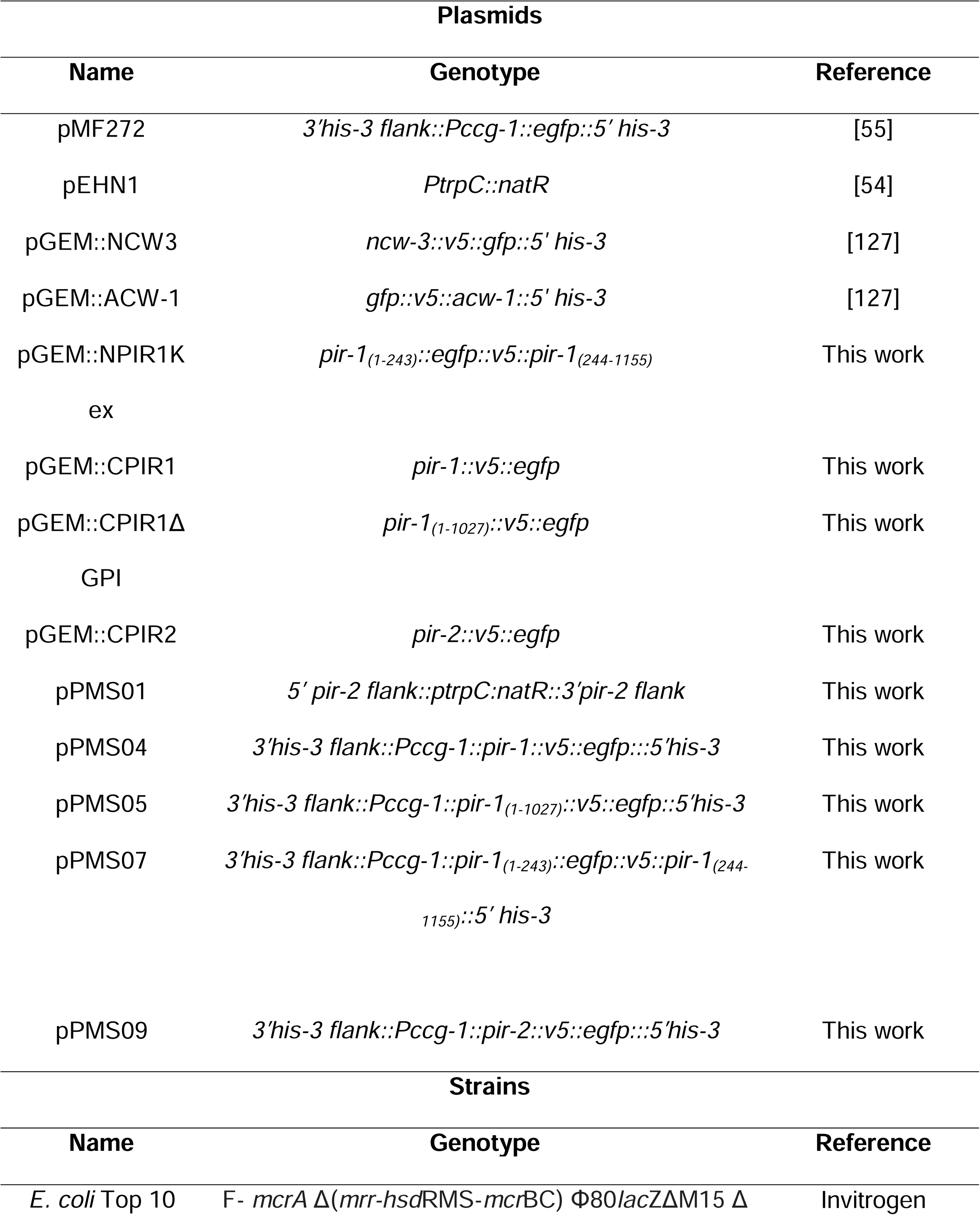
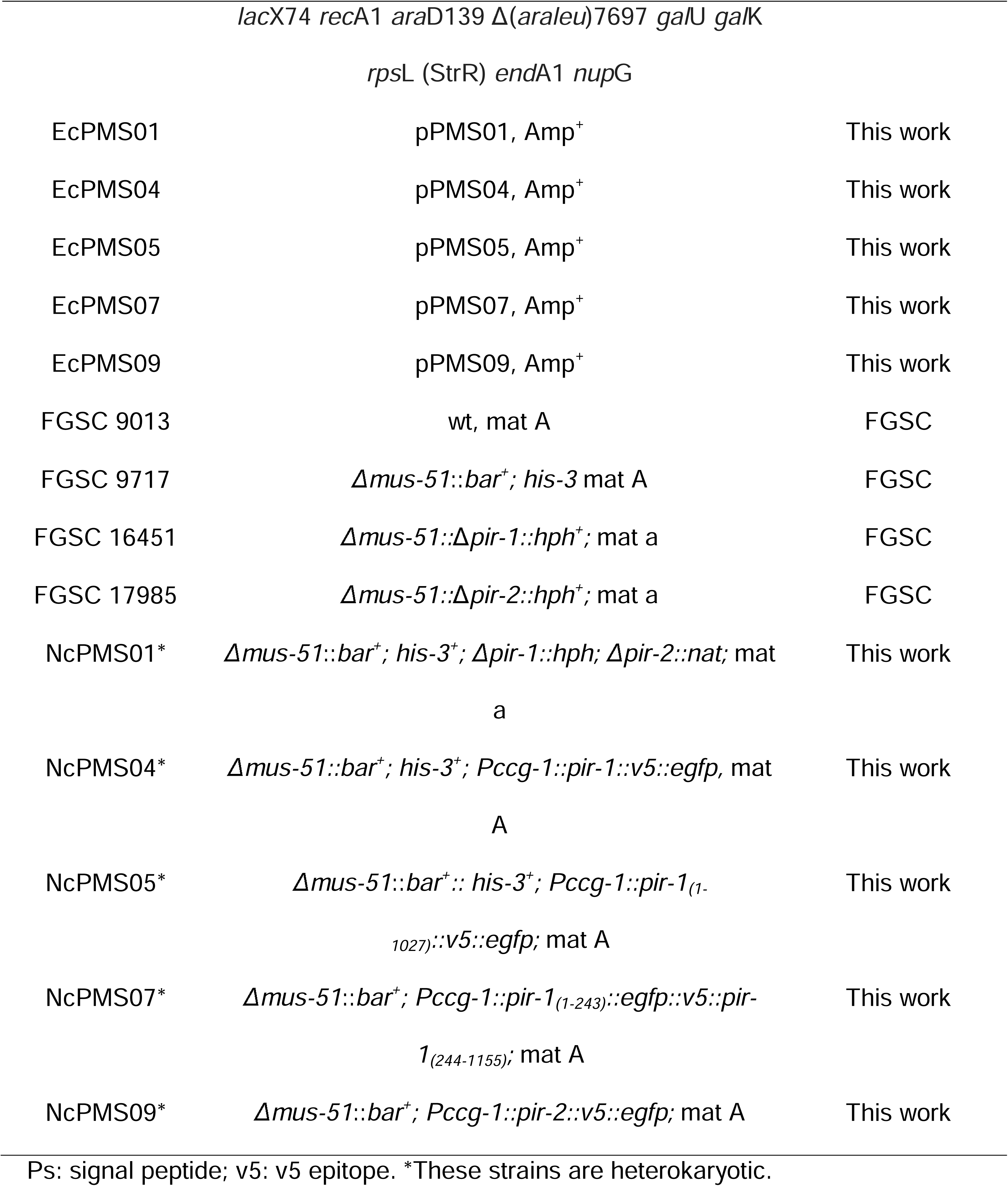
Plasmids and strains used in this work.

*N. crassa* strains were cultivated at 30 °C on Vogel’s minimal medium (VMM) agar [52], supplemented with 1.5 % (w/v) sucrose and, if necessary, 0.5 mg/mL histidine or 100 µg/mL nourseothricin. Fresh conidia were electroporated in a Bio-Rad Gene Pulser Xcell Total System under the following conditions: 600 Ohms, 25 µFD and 1.5 kV. After the electric pulse, conidia were recovered in 1 mL 1 M sorbitol, incubated at 30 °C for two hours with gentle agitation, and plated on FGS agar [53] for five days at 30 °C.

### 2.3 Molecular techniques

#### N. crassa double knock-out construction

To generate the double knockout N. crassa Δpir-1/Δpir-2 strain, 5’ pir-2 flank::PtrpC::nat::3’ pir-2 flank assembly was constructed using NEBuilder Assembly Kit (New England Biolabs). Each assembly fragment was either synthesized (T4 Oligo, Irapuato, Mexico) or PCR amplified using Platinum High Fidelity Taq polymerase (Thermo Fisher) and then gel purified (GenElute DNA Extraction Kit, Sigma). PCR primers were designed to provide each amplicon an ending homologous to its adjacent fragments and make possible the assembly. In addition, two melting temperatures (Tm) were considered for each primer: a total Tm and a partial one (Table S1), where partial Tm must be around 50 °C in all cases and corresponded to the fragments overlapping zone. 5’-pir-2 and 3’- pir-2 flanks were PCR amplified from N. crassa FGSC 9013 (WT) genomic DNA with 2PIR 2-1, 2PIR 2-2, and 2PIR 2-5 and 2PIR 2-6 primers (Table S1), respectively. natR and its corresponding promoter (PtrpC) were amplified from pEHN1 [54] with 2PIR 2-3 and 2PIR 2-4 primers. Three fragments were assembled in a one step assembly reaction and reamplified by PCR, and then ligated into pGEM T-Easy Vector (PROMEGA). The resulting plasmid (pPMS01) was then cloned in E. coli TOP10, and purified by miniprep (GenElute Plasmid Miniprep Kit, Sigma). To discard any mutation, the plasmid was Sanger sequenced (LANGEBIO-CINVESTAV, Irapuato, Mexico).

#### N. crassa his-3-targeted assemblies

Three assemblies were built for expression of co-translational fusions of PIR-1 with eGFP: *pir-1_1-243_::egfp::v5::pir-1_244-1155_*(eGFP-PIR-1Kex)*, pir-1::v5::egfp* (PIR-1-eGFP), and *pir-1*_1-1027_*::v5::egfp* (PIR-1ΔGPI-eGFP); and one assembly for expression of PIR-2 fusion with eGFP: *pir-2::v5::egfp* (PIR-2-eGFP). A detailed procedure for each assembly construction is provided in the supplementary material and Figure S1. Once assemblies were obtained, they were double digested with XbaI and BstBI (Anza, ThermoFisher), and ligated into pMF272 [55] to put the expression cassette under the control of the *Pccg-1* promoter, and add *his-3* and *his-3* 3’ flank regions which target them to the *his-3* locus of *N. crassa* genome. The latter led to plasmids pPMS04, pPMS05, pPMS07, and pPMS09, which were cloned in *E. coli Top 10* leading to EcPMS04, EcPMS05, EcPMS07 and EcPMS09 strains, respectively. Plasmids were characterized by Sanger sequencing.

#### N. crassa transformation and selection

*N. crassa* transformation was performed as described above. Circular plasmids were directed either to the *his-3* locus of *N. crassa* FGSC 9717 for overexpressing strains, or *pir-2* locus of *N. crassa* FGSC 16451 (Δ*pir-1)* for double Δ*pir-1*/Δ*pir-2* knockout strain. Transformants were selected either for nourseothricin resistance (NcPMS01), or histidine prototrophy (NcPMS04, NcPMS05, NcPMS07 and NcPMS09). *N. crassa* heterokaryotic transformants were molecularly confirmed by genomic PCR (Figure S2 and S3). For a detailed procedure see the supplementary material.

### 2.4 Phenotypic analysis

#### Morphology

*N. crassa strains* WT, Δ*pir-1*, Δ*pir-2*, and Δ*pir-1/*Δ*pir-2* were phenotypically analyzed by testing morphological differences in VMM culture media. *Growth rate*. 1×10^6^ conidia were inoculated in VMM agar plates and incubated at 30 °C overnight. Then, 1 cm^2^ colonized agar was transferred into a fresh VMM plate and incubated again at 30 °C. After four hours, once the growth rate was stabilized, growth was measured every two hours for 16 h. The assay was performed in quintuplicate (n = 5). *Branching patterns*. To estimate branching patterns, lateral branches were quantified in parent hyphae from the closest branch to the tip and up to 500 μm apart from it (n = 20). Branching was reported as number of branches (#br)/500 μm. For statistical analysis methods see section 2.7.

### 2.5 Stress assays

#### Thermal stress assays

WT and mutant *Neurospora* strains (Δ*pir-1,* Δ*pir-2, and* Δ*pir-1/*Δ*pir-2*) were tested for thermal sensitivity. 1×10^6^ conidia were cultivated on VMM agar at 30 °C overnight (16 h); then, 1 cm^2^ colonized agar was transferred into a fresh VMM plate, and further incubated at 45 °C. Growth rate was measured every two hours during 8 hours after the thermal shift. The assay was performed in quintuplicate (n = 5).

#### Calcofluor white (CFW) susceptibility assays

First, 5×10^3^ conidia were cultivated on VMM agar at 30 °C; then, 1 cm^2^ colonized agar was transferred into a fresh VMM plate supplemented with 1 mg/ml CFW and VMM without CFW was used as control. Growth rate was measured every two hours during 16 h after 12 h culture transfer to CFW plates. The assay was performed in duplicate (n=2). For statistical analysis methods see section 2.7.

#### Osmotic stress assay

5×10^3^ conidia were cultivated on VMM agar at 30 °C overnight (16 h); then, 1 cm^2^ colonized agar was transferred into a fresh VMM plate supplemented with 4% (w/v) NaCl and VMM without NaCl was used as control. Growth rate was measured every two hours during eight hours after 12 h culture transfer to NaCl plates. The assay was performed in triplicate (n=3). For statistical analysis methods see section 2.7.

### 2.6 Live imaging

Confocal images were collected using an OlympusFluoview FV-1000 inverted confocal microscope at the Laboratorio Nacional de Microscopía Avanzada (LNMA), CICESE, Ensenada, Mexico. All samples were observed using the inverted agar technique [56] with the apochromatic objective 60X/1.40 and immersion oil. A Xenon laser was used for eGFP excitation at 488 nm, and emission was collected at 505-605 nm wavelength. All images were analyzed with FIJI [57].

### 2.7 Statistical analysis

A One-way analysis of variance (ANOVA) was conducted to compare *N. crassa* mutant strains (Δ*pir-1*, Δ*pir-2*, Δ*pir-1/*Δ*pir-2*) and the control, *N. crassa* WT. Fisher LSD test was performed. Statistical significance was determined at p<0.05. All statistical analyses were performed in Rstudio.

## 3 RESULTS

### 3.1 PIR proteins are exclusive to Ascomycota

To explore the diversity of PIRs beyond the already characterized PIRs from yeasts [11,30,58], the yeast PIR-domain (PF00399) was used as a query to explore the diversity of PIRs in the tree of life. Despite systematic searches in different databases (Ensembl All, UniProtKB, and SwissProt), only sequences from fungi were retrieved. To assess the distribution of PIRs among fungi, 840 fungal reference proteomes from the NCBI refseq database (October, 2022) were downloaded and scanned against the yeast PIR-domain. A total of 284 species were positive to at least one PIR domain; in order to prevent false positives, those sequences were analyzed to identify the five amino acids (Q, D, G, Q, Q) biochemically essential for anchoring the PIR domain to the wall [19], which led to a final set of 225 species (Figure 1).

**Figure 1.**
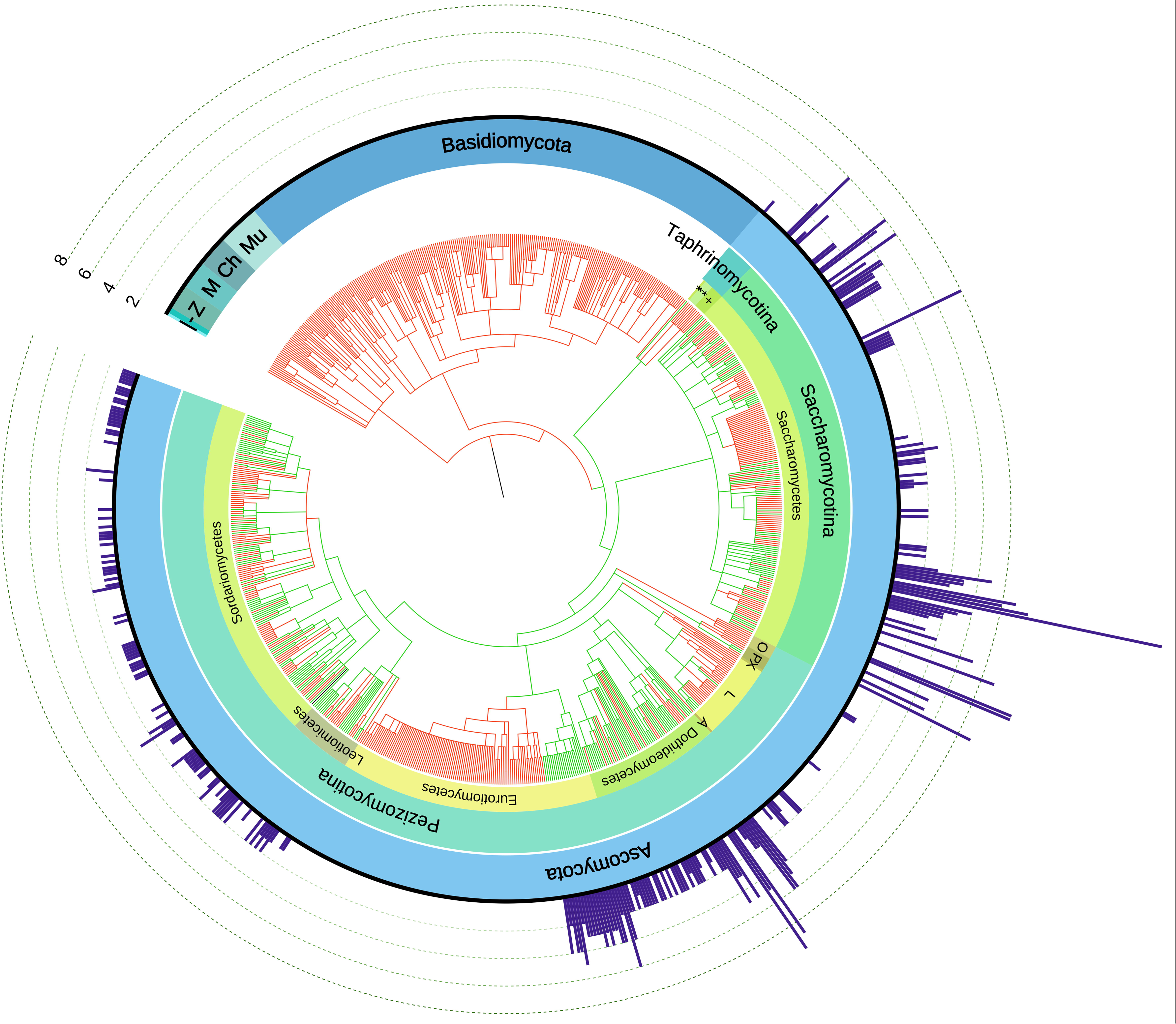
Allocation of PIRs in the fungal tree of life. PIR-domains containing proteins were retrieved from the proteomes of 840 fungal reference species. Within the species phylogeny, green colored branches indicate a species with at least one PIR domain-containing protein. Fungal classes are indicated in the innermost label circle, while subdivision and phylum are indicated in progressive outer ones. Projected radial bars indicate the number of PIRs encoded in the genome of each species. The fungal phylogeny was taken from [49] and the presence of PIRs in fungal families was annotated with iTOL [128]. |, Cryptomycota; -, Blastocladiomycota; Z, Zoopagomycota; M, Microsporidia; Ch, Chytridiomycota; Mu, Mucoromycota; **, Schizosaccharomycetes; *, Taphrinomycetes; +, Pneumocystidiomycetes; O, Orbiliomycetes; P, Pezizomycetes; X, Xylonomycetes; L, Lecanoromycetes; A, Arthoniomycetes.

All positive species were exclusive to Ascomycota (Figure 1, Table S2). They were allocated in all three Ascomycota subdivisions (Taphrinomycotina, Saccharomycotina, and Pezizomycotina) but in only eight classes: Taphrinomycetes (1 species), Saccharomycetes (63)—both early divergent ascomycetes—, and also in Pezizomycetes (2), Lecanoromycetes (5), Dothideomycetes (43), Eurotiomycetes (23), Leotiomycetes (15), and Sordariomycetes (52) (Figure 1). PIRs in Ascomycota classes were unevenly distributed among their species. Within Saccharomycetes, discrete PIRs containing clusters were clearly observed at the genus level only among *Ogataea*, *Brettanomyces*, *Clavispora*, *Kluyveromyces*, *Naumovozyma*, *Tetrapisispora*, and *Eremothecium*. Within Dothideomycetes, unlike Saccharomycetes, Leotiomycetes, and Sordariomycetes, most species contained PIRs. On the other hand, within Eurotiomycetes, a monophyletic group comprising the major clades Eurotiomycetidae and Chaetothyriomycetidae, PIRs were exclusive to Chaetothyriomycetidae, appearing in all its genera. Moreover, PIRs were conspicuously more abundant in species of Saccharomycetes (4.25 proteins/species on average), followed by species of Eurotiomycetes (3.40 proteins/species on average), Dothideomycetes (2.74 proteins/species on average), and lastly, species of Leotiomycetes and Sordariomycetes contained only one single PIR on average (Figure 1).

### 3.2 PIRs group in 11 functional clusters mainly defined by the allocation of PIR domains within the protein

To identify sub-groups among proteins containing PIR domains (Figure 2A), a sequence similarity network (SSN) analysis was carried out [48]. EFI-EST analysis resulted in 11 clusters, five of which were two-sequence clusters, and 22 singletons that were discarded for downstream analysis. Each cluster was analyzed for features relevant to PIRs and CWPs, including secretion (SP) and GPI signals (GPI) [41,42], the absence of transmembrane domains (TM) [59], cysteine-rich domains [34], serine/threonine composition [60], functional domains [39], and intrinsic disorder [45]. The graphical representation of the six main PIR clusters is shown in Figure 2B and their characteristics are summarized in Table S2.

**Figure 2.**
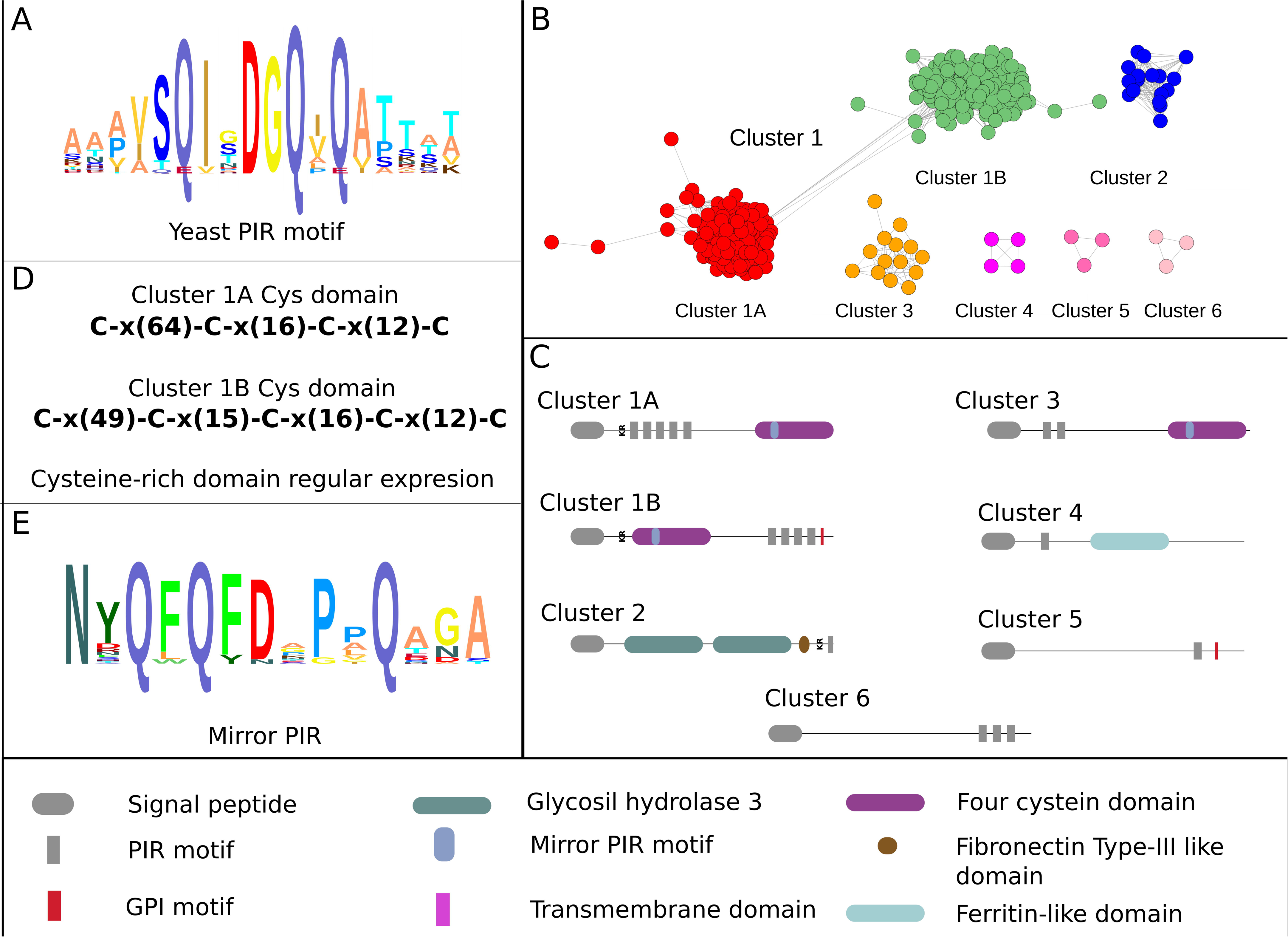
Diversity of fungal PIRs. (A) The PIR domain is well conserved all over the PIRs universe. (B) Retrieved fungal PIRs were clustered by a sequence similarity network (SSN) analysis using EFI-EST [48]. Eleven clusters were identified, but only the six more populated were analyzed. (C) Each cluster was defined by a particular architecture where PIR domains could be located either close to the N-terminus, the middle or the C-terminus of the protein. Major clusters 1A and 1B were the only ones that contained both PIR domains and a Cys-rich domain. (D) Cys-rich domain is defined by four Cys residues among yeasts PIRs (cluster 1A), while among filamentous fungi PIRs (cluster 1B) the Cys-rich domain is defined by five Cys-residues. (E) Within the Cys-rich domain of PIRs from clusters 1A and 1B an inverted PIR domain, named mirror PIR, was found.

The vast majority of PIRs were grouped into one main cluster (cluster 1, 82% of total PIRs); however, two independent, but interconnected sub-clusters were clearly observed and named cluster 1A (45% of total PIRs) and cluster 1B (36.75% of total PIRs) (Figure 2B). Cluster 1A contained PIRs from yeast species with the previously reported canonical PIRs architecture (N-terminal PIR domains, C-terminal Cys-rich domain) (Figure 2C), while cluster 1B contained exclusively proteins from filamentous fungi that, unlike cluster 1A PIRs, showed an inverted architecture (N-terminal Cys-rich domain, C-terminal PIR domains). Clusters 2 to 6 were less populated and also contained PIRs from filamentous fungi (Table S2).

Up to 12 PIR domains in a single PIR were found distributed throughout the entire sequence (Figure 2C). In clusters 1A, 3, and 4, PIR domains were located close to the N-terminus; while in clusters 1B, 2, 5, and 6, PIR domains were placed close to the C-terminus of the protein. A multiple sequence alignment revealed four (C_1_-x(64)-C_2_-x(16)-C_3_-x(12)-C_4_) and six (C_-1_-x(27)-C_1_-x(49)-C_x_-x(15)-C_2_-x(16)-C_3_-x(12)-C_4_) Cys-rich domains in clusters 1A and 1B, respectively (Figure 2D, S4). They were consistently located at the C-terminus among yeasts PIRs (cluster 1A), but at the N-terminus among filamentous fungi PIRs (cluster 1B) (Figure 2C, S4). In PIRs cluster 1B, the first Cys (C_-1_) of the domain was upstream of the Kex2 processing signal (KK/KR); thus, the regular expression of the mature PIR protein was a five Cys-rich domain, C_1_- x(49)-C_x_-x(15)-C_2_-x(16)-C_3_-x(12)-C_4_. The sequence alignment also revealed that the additional Cys (C_x_) of the domain was an insertion between Cys-1 and Cys-2 of the canonical yeasts Cys-rich domain (Figure S4). Detailed sequence analysis also showed that a subset of cluster 1B contained only a three Cys domain. Unlike cluster 1B, cluster 3 Cys-rich domain was expanded with an insertion of variable length (Figure S5), from only an additional Cys (*Exophiala xenobiotica:* XP_013320788) to 46 additional amino acids (*Fonsecaea erecta*: XP_018698287), revealing a domain expansion of up to 47 amino acids between Cys-1 and Cys-2 with a regular expression C_1_-x(49)-C_x_-x([15–47])-C_2_-x(16)-C_3_-x(12)-C_4_ (Figure S5). Noteworthy, clusters 2, 4, 5, and 6 lacked a Cys-rich domain in the PIRs (Figure 2C, Table S2). In addition, between Cys-1 and Cys-2 of the Cys-rich domains in clusters 1A and 1B, we identified a highly conserved region with an inverted PIR domain, which we have named mirror-PIR (Figure 2E). The three glutamine and aspartic acid residues required for the cell wall anchoring were highly conserved in both PIR and mirror-PIR domains (Figures 2A, E and S4). However, in cluster 1A, between the second and third glutamine, mirror-PIR additionally had three highly conserved proline residues, while only one proline was conserved in cluster 1B mirror-PIR. On average, mirror-PIR was five amino acids longer than direct PIR domains (Figures 2A, E and S4). On the other hand, cluster α and cluster β, formally non PIRs since they lacked PIR domains, showed a typical Cys-rich domain with a mirror-PIR domain within (Table S2).

In addition to the definitory PIR domain, PIRs showed a high diversity of motifs and domains distributed along their entire sequence. Consistently, filamentous fungi PIRs had a GPI modification signal to the C-terminus that was completely absent in yeasts PIRs. None of the PIRs analyzed contained transmembrane domains. All members of the main PIR clusters (1A and 1B), had Kex2 processing signals, but 20 sequences in 20 species lacked signal peptide (Table S2). Proteins in cluster 2 harbored a family 3 glycosyl hydrolase domain (GH3; PF01915.24/25), kept the Kex2 processing signals located at the C-terminus (Figure 2C), and were the only ones with catalytic domains. Cluster 4 harbored a Ferritin_2-like domain (PF00399.22). A full description of PIRs clusters is provided in the supplementary Table S2. Overall, all ascomycetous species were predicted to have at least one PIR with one PIR domain along their sequence. They had at least one protein belonging to cluster 1A, except for *Neurospora tetrasperma*. Species harboring 2 predicted PIRs belong to clusters 2, 3, 4, and 5, and can be considered variants of the canonical filamentous fungi PIRs architecture (Figure 2C).

### 3.3 PIRs are intrinsically disordered

Since protein repeats, characteristic of CWPs, have been associated with intrinsic disorder [61,62], a common way to form intermolecular complexes [63,64] and a necessary asset to secrete CWPs through the cell wall in bacteria [65], intrinsic disorder was assessed in identified PIRs (Table S2). In general, filamentous fungi PIRs (cluster 1B) were more intrinsically disordered (93.03% of total proteins had at least 1 intrinsically disordered region, IDR, longer than 50 amino acids; 99.31 % of total proteins had at least 1 IDR longer than 30 amino acids), than cluster 1A yeasts PIRs (77.6 % of total proteins had at least 1 IDR longer than 50 amino acids; 91.26 % of total proteins had at least 1 IDR longer than 30 amino acids). In less populated PIRs clusters, IDRs longer than 50 amino acids appeared in 100% members of clusters 2, 3, 5, and 6; in cluster 4, 50% members have at least one IDR longer than 50 amino acids. On the other hand, disordered regions longer than 30 amino acids were observed in 100% members of clusters 2, 3, 4, 5, and 6 (Table S2).

In order to assess whether intrinsic disorder was directly associated to PIR domains, one representative PIR from each cluster was taken and intrinsic disorder inferences from three different predictors (FIDPnn, IUPRED3 and ESpritz) were matched with PIR domains distribution (Figure S6). Although there was a match between PIR motifs and intrinsic disorder in several PIRs, it was not a general trend. Nevertheless, in cluster 1B, where PIR domains are located at the C-terminus, intrinsic disorder was consistently found in the same region (Figure S6, Table S2). The same analysis was performed for well studied classical PIRs from *S. cerevisiae* (PIR1p, PIR2p, PIR3p, and PIR4p) (Figure 3A, B, C, and D) and *N. crassa* PIR-1 and PIR-2, two PIRs with an inverted architecture typical of filamentous fungi (Figure 3E and F). Except for *Sc* PIR4p (Figure 3D), all the other analyzed PIRs contained long disordered regions (Figure 3). Only the totality of PIR domains of *Sc* PIR2p and *Nc* PIR-2 were within a disordered region; half of the PIR domains of *Sc* PIR1p, *Sc* PIR3p and *Nc* PIR-1 were under a disordered region, while all PIR domains of *Sc* PIR4p were in ordered regions. AlphaFold prediction of PIRs tertiary structures confirmed disorder prediction. All proteins contained a well structured core (around the Cys-rich domain in the case of PIRs that contained it, *Sc* PIR1p, PIR2p, PIR3p, PIR4p, and *Nc* PIR-1) surrounded by two disordered arms (Figure 3). It is noticeable that PIR domains in IDRs are, nevertheless, prone to form β-hairpins (Figure S7).

**Figure 3.**
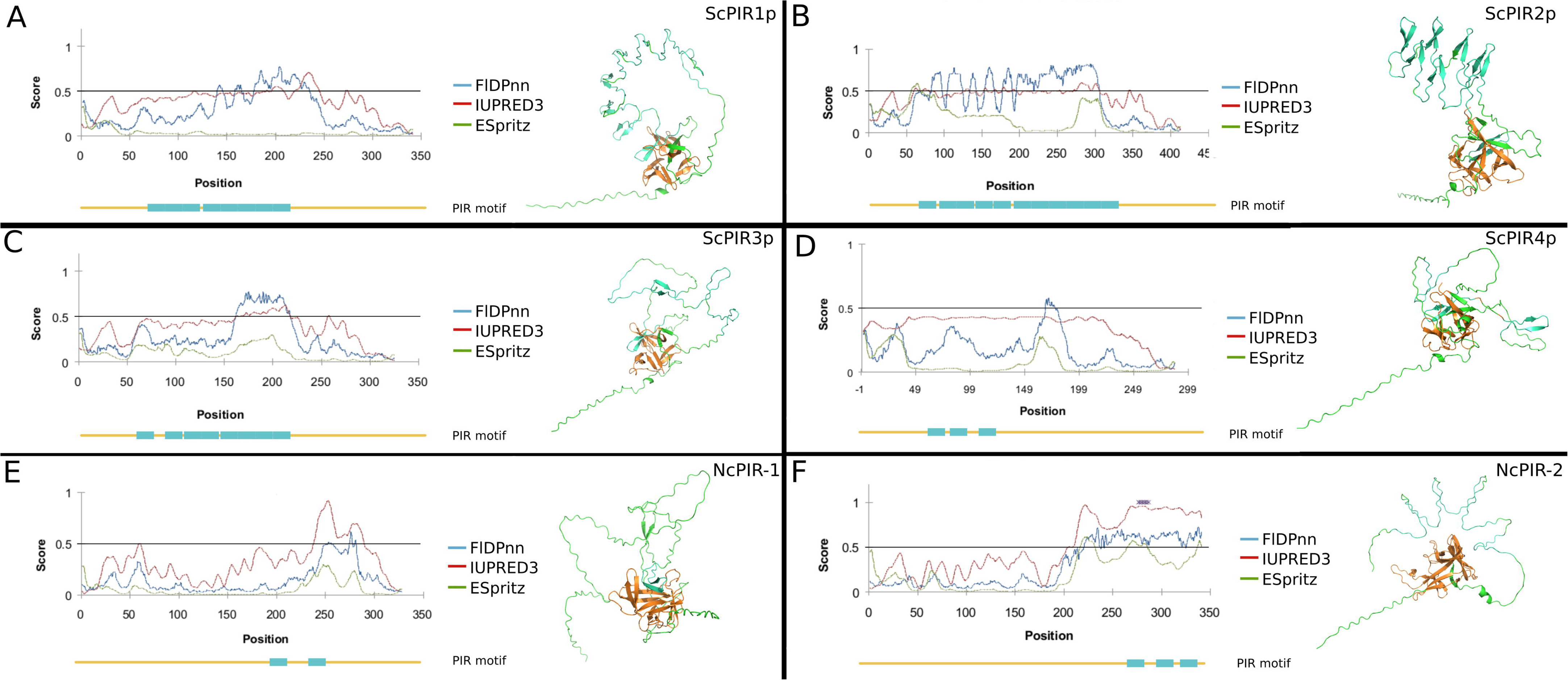
Intrinsic disorder is not linked to PIR domains. Intrinsic disorder was estimated with FIDPnn (blue line), IUPRED3 (red line) and ESpritz (green line) for (A) ScPIR1p, (B) ScPIR2p, (C) ScPIR3p, (D) ScPIR4p, (E) NcPIR-1 and (F) NcPIR-2, and matched with the allocation of PIR domains within each protein (lower line, turquoise blocks). In addition, the structural model of each analyzed PIR was predicted with AlphaFold [50]. Consistently, PIRs contained a well structured core (ochre) surrounded by two intrinsically disordered arms (green).

### 3.4 Single deletion of *N. crassa pir-1* and *pir-2* did not compromise the cell wall stability, but the double mutant did

Putative PIRs among filamentous ascomycetes were previously inferred for *N. crassa*, *Magnaporte grisea, Gibberella zeae,* and *Blumeria graminis* [34]. *N. crassa* harbors 2 PIRs from different clusters: PIR-1 belongs to cluster 1B (Table S2); it contains a Cys-rich domain close to the N-terminus and two PIR domains close to the C-terminus. On the other hand, PIR-2 belongs to cluster 6 (Table S2); it is devoid of a Cys-rich domain, but contains three PIR domains. Because of their inverted architecture relative to canonical yeasts PIRs and the presence of additional post-translational signals that could modify their physiological role (i.e. GPI), PIR-1 and PIR-2 were further characterized as models of filamentous fungi PIRs.

To functionally characterize PIR-1 and PIR-2, homokaryotic single gene-deletion mutant *N. crassa* (Nc) strains Δ*pir-1* (FGSC16451) and Δ*pir-2* (FGSC17985), as well as the heterokaryotic double-gene deletion mutant Δ*pir1/*Δ*pir-2* (NcPMS01) were analyzed. Nc Δ*pir-1* and Nc Δ*pir-2* strains were acquired from the FGSC and molecularly confirmed by PCR (Figure S2A and B). Nc Δ*pir1/*Δ*pir-2* double mutant was constructed from Nc Δ*pir-1*, which was also molecularly confirmed by PCR (Figure S2C). Despite numerous efforts, this strain was analyzed heterokaryotic since it was not possible to isolate a homokaryotic one.

Compared to Nc *WT*, Δ*pir-1* and Δ*pir-2* single mutant strains exhibited no morphological alterations showing regular dichotomic ramifications (Figure 4A-C). However, colony density of Nc Δ*pir-1/*Δ*pir-2* was negatively affected (Figure 4D). Growth rate of Nc WT (4.10±0.53 mm/h) was not statistically different (p>0.05) to that of Nc Δ*pir-1* (4.33±0.07 mm/h) and Nc Δ*pir-2* (3.67±0.32 mm/h). However, there were significant differences between Nc Δ*pir-1* and Nc Δ*pir-2*; Nc Δ*pir-1* grew faster. Nc Δ*pir-1/*Δ*pir-2* growth rate was negatively affected (2.08±0.44 mm/h), showing only 50% the growth rate of Nc WT, Δ*pir-1* and Δ*pir-2* strains (Figure 4E). All analyzed mutant strains revealed different ramification patterns (Figure 4F). Branching of Nc Δ*pir-1,* Nc Δ*pir-2* and Nc Δ*pir-1*/Δ*pir-2* were statistically different (p<0.05) to that of Nc WT. However, whereas Nc Δ*pir-1* showed a decreased number of branches (2.45±0.88 branches/500 μm) relative to Nc WT (3.05±0.82 branches/500 μm), in Nc Δ*pir-2* branching increased (3.80±1.05 branches/500 μm). On the other hand, Nc Δ*pir-1*/Δ*pir-2* branched (2.30±0.65 branches/500 μm) similarly to Nc Δ*pir-1* (Figure 4F).

**Figure 4.**
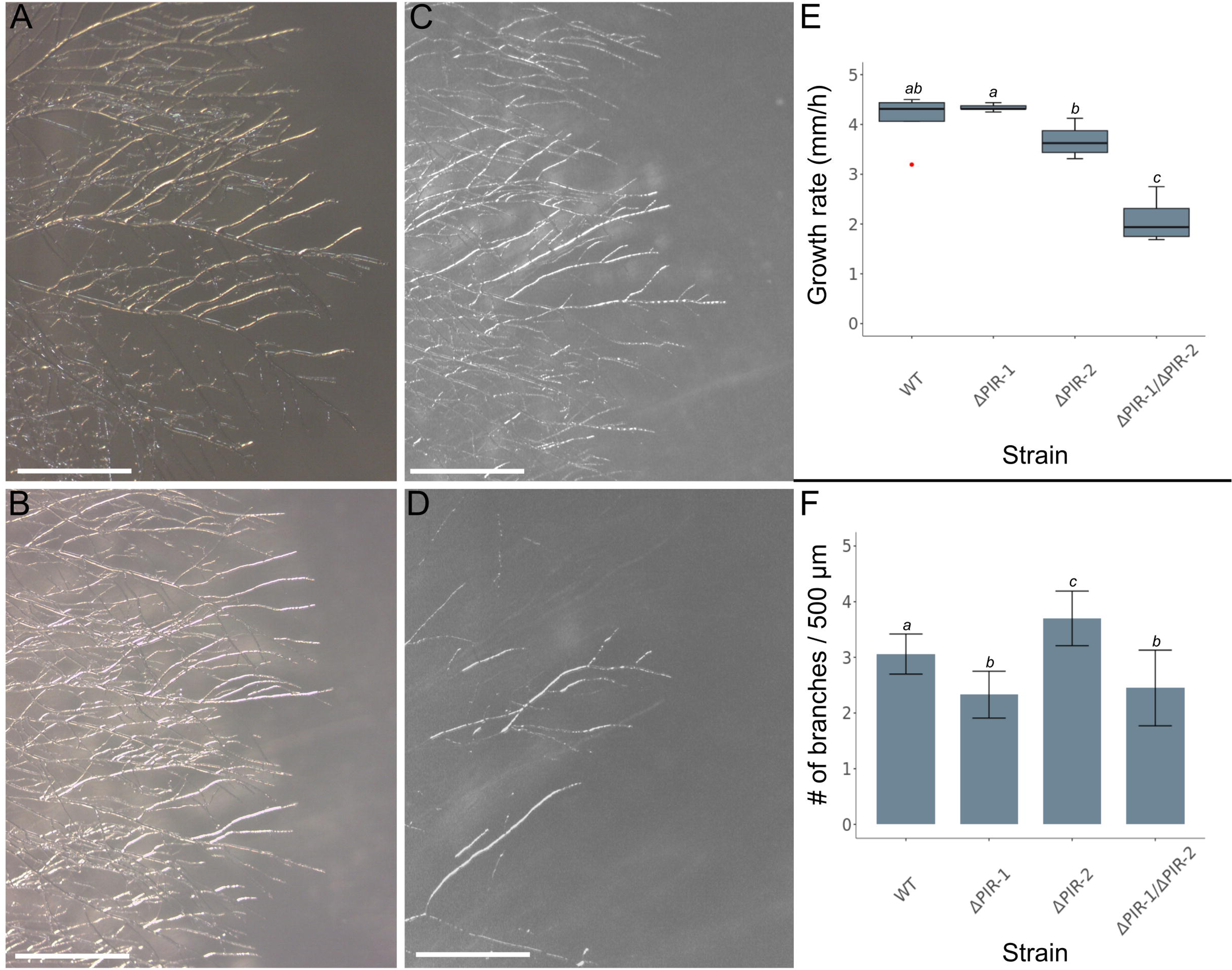
*N. crassa* Δ*pir-1*/Δ*pir-2* double mutation synergistically affects the hyphal growth rate. Nc Δ*pir-1* (FGSC16451) and Nc Δ*pir-2* (FGSC17985) mutants, as well as the double knockout Nc Δ*pir-1/*Δ*pir-2* strains (NcPMS01) were studied to determine their cellular role. (A-C) Microscopy analysis of Nc WT, Δ*pir-1*, and Δ*pir-2* showed no major morphological differences between them; however, (D) colony density of Nc Δ*pir-1/*Δ*pir-2* decreased. Images are representative of 5 fields; scale bar, 20 μm. (E) Growth rates were not significantly different between Nc Δ*pir-1 and* Nc Δ*pir-2* relative to Nc WT, which suggested no functional compromise. Nevertheless, Nc Δ*pir-1/*Δ*pir-2* growth rate was significantly slower than that of the other analyzed *N. crassa* strains. Growth rate measurements were the average of 5 replicates. (F) A statistically significant difference in branching frequency between Nc WT, Δ*pir-1* and Δ*pir-2* was observed. However, branching of double knockout Nc Δ*pir-1/*Δ*pir-2 was the same as* Nc Δ*pir-1*. Branching frequency was estimated with an n=20. A one-way analysis of variance (ANOVA) was conducted to compare mutant strains in each measurement. Fisher LSD test was performed. Statistical significance was determined at p<0.05.

To directly test whether Nc Δ*pir-1* and Δ*pir-2* knockouts caused a cell wall defect, a cell wall integrity assay was performed with calcofluor white (CFW), an inhibitor of chitin synthesis [66–68]. After testing different CFW concentrations, 1 mg/ml CFW was set as the inhibitory concentration to perform the assay. Under this stress condition, the growth rate of Nc WT, Nc Δ*pir-1* and Nc Δ*pir-2* were similarly reduced 16.09±6.7% (0.07 mm/h), 17.0±7.2% (0.06 mm/h), and 10.6±2.6% (0.03 mm/h) regarding themselves grown in absence of the stressor (Figure 5A). Nevertheless, Nc Δ*pir-1/*Δ*pir-2* showed a growth rate reduction of 46.93±1.6% (0.11 mm/h) (Figure 5A). Although single mutants were not significantly stressed by CFW, which indicated no cell wall defects, the magnitude of Nc Δ*pir-1/*Δ*pir-2* growth rate reduction in presence of CFW suggested a synergic effect of both mutations that led to a major cell wall defect.

**Figure 5.**
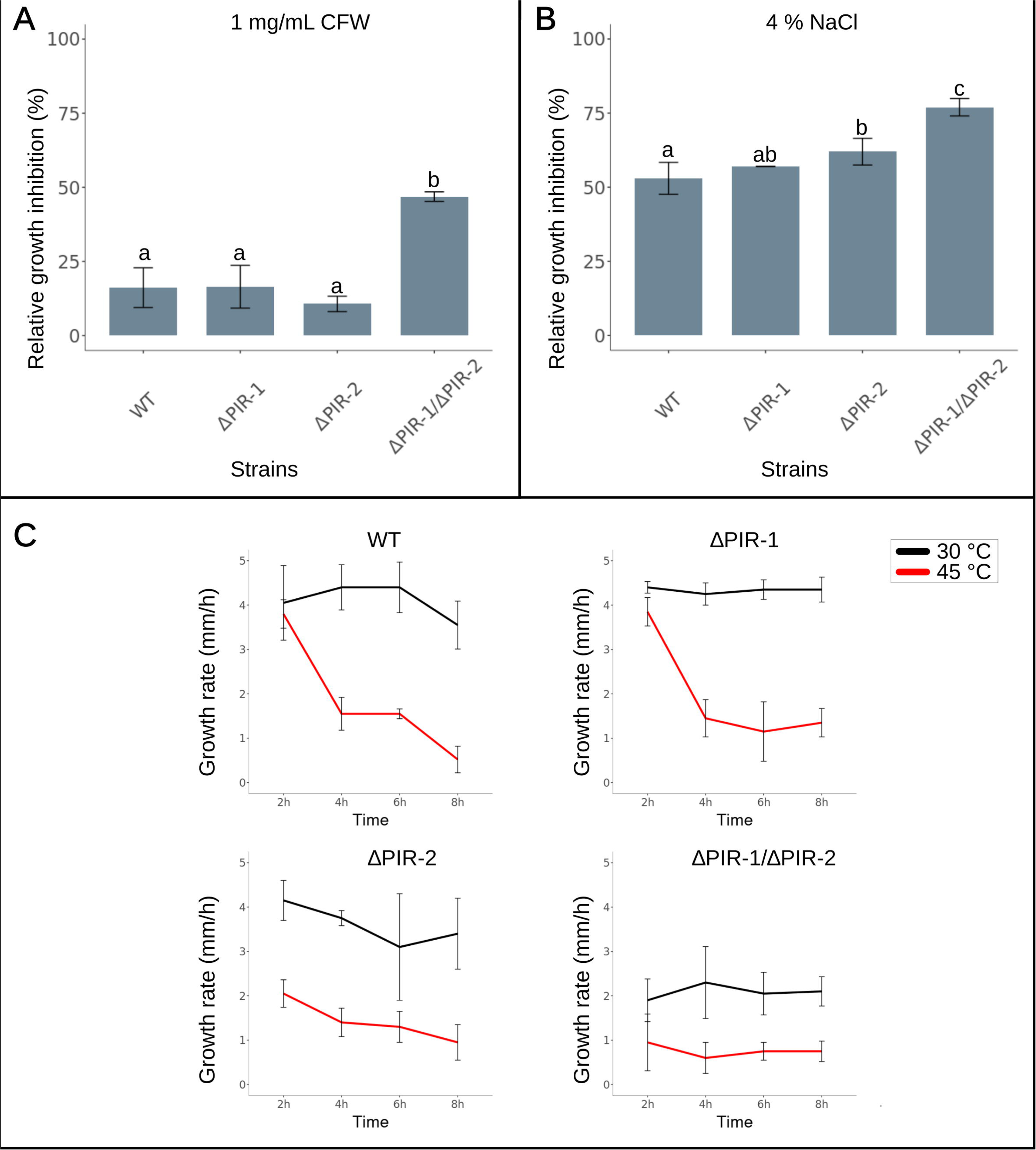
PIR-1 and PIR-2 act together to maintain the cell wall stability. (A) Nc *WT* (FGSC 9103), Nc Δ*pir-1* (FGSC 16451), Nc Δ*pir-2* (FGSC 17985), and Nc Δ*pir-1*/Δ*pir-2* (NcPMS01) were grown in (A) 1 mg/ml CFW, (B) 4% NaCl, and (C) 30 and 40 °C to assess any cell wall defect that compromises its stability. (A) Both Nc Δ*pir-1 and* Δ*pir-2* were similarly sensitive to CFW as Nc WT indicating no cell wall defect due to the absence of any of both proteins. Nevertheless, Nc Δ*pir-1*/Δ*pir-2* double mutant was sensitive to CFW in a magnitude that suggested a synergic effect. The average of 2 replicates of each strain grown in 1 mg/ml CFW relative to each strain cultivated in absence of the stressor is shown. (B) Unlike CFW assays, 4% NaCl osmotic stress had a progressive inhibitory effect on Nc *WT*, Nc Δ*pir-1,* Nc Δ*pir-2,* and Nc Δ*pir-1*/Δ*pir-2*. Despite the magnitude of the osmotic stress effect on Nc Δ*pir-1*/Δ*pir-2* was stronger than that observed for the single mutants, it was neither additive nor synergic. The average of 3 replicates of each strain cultivated on 4% NaCl relative to each strain cultivated in absence of the stressor is shown. (C) All strains under analysis were grown at 30 °C and 45 °C for 8h, and their growth rate determined. At 45 °C, Nc *WT* and Nc Δ*pir-1* stepwisely decreased their growth rate along the incubation time. In contrast, Nc Δ*pir-2* showed a reduced growth rate from the beginning of the incubation at 45 °C, while the double mutant Nc Δ*pir-1*/Δ*pir-2* showed a drastic growth rate decrease at both 30 °C and 45 °C. The average growth rate of 5 replicates of each strain cultivated at 30 °C or 45 °C is shown. A one-way analysis of variance (ANOVA) was conducted to compare mutant strains in each stress assay. Fisher LSD test was performed. Statistical significance was determined at p<0.05.

Because of the detected cell wall damage in Nc Δ*pir-1*/Δ*pir-2* mutant, as well as the report of the relationship of PIRs with the osmotic stress response [17], growth of Nc WT, Nc Δ*pir-1,* Nc Δ*pir-2,* and Nc Δ*pir-1*/Δ*pir-2* in 4% NaCl was assessed (Figure 5B). Although the growth rate of Nc WT was reduced by 53±5.39% (0.23 mm/h), the growth rate of Nc Δ*pir-1* (57±0%; 0.22 mm/h reduction), Nc Δ*pir-2* (62±4.51%; 0.21 mm/h reduction), and Nc Δ*pir-1*/Δ*pir-2* (77±2.95%; 0.17 mm/h reduction) was increasingly affected regarding their own growth rate in absence of the stressor (Figure 5B). Unlike the case of CFW inhibition assay, the effect of the osmotic stress on Nc Δ*pir-1*/Δ*pir-2* mutant was not the addition of each single mutation effect, neither synergic. To further study the biological role of PIR-1 and PIR-2, the growth rate of Nc WT, Nc Δ*pir-1,* Nc Δ*pir-2,* and Nc Δ*pir-1*Δ*pir-2 at 30* °C and 45 °C was measured (Figure 5C). At 30 °C, after 8 h cultivation, no significant growth difference was observed between Nc *WT* (4.05±0.84 mm/h), Nc Δ*pir-1* (4.4±0.13 mm/h) and Nc Δ*pir-2* (3.67±0.32 mm/h) (Figure 5C); however, the double mutant Nc Δ*pir-1/*Δ*pir-2* (1.9±0.48 mm/h) showed a growth rate decrease of around 50% along all the tested time (Figure 5C). At 45 °C, the restrictive temperature, both Nc *WT* and Nc Δ*pir-1* showed a similar growth behavior; after 2 h incubation at 45 °C, their growth rate (3.8±0.23 mm/h and 3.85±0.32 mm/h, respectively) was almost the same as 30 °C (4.05±0.84 and 4.4±0.13 mm/h, respectively) (Figure 5C). Nevertheless, after 4 h incubation at 45 °C their growth rate drastically decreased to 1.55±0.23 mm/h (Nc *WT*) and 1.45±0.42 (Nc Δ*pir-1*), which kept close to this value after 8 h incubation at the restrictive temperature (0.52±0.35 mm/h and 1.35±0.32 mm/h, respectively) (Figure 5C). In contrast, Nc Δ*pir-2* showed a decreased growth rate at 45 °C from the beginning of the incubation (2.05±0.31 mm/h) and continuously decreased (0.95±0.40 mm/h) till 8 h incubation (Figure 5C). The double heterokaryotic mutant Nc Δ*pir-1*/Δ*pir-2* showed an even lower growth rate from the beginning of the incubation at 45 °C (1.18±0.5 mm/h), which was approximately constant (0.75±0.23 mm/h) till the end of the tested time (Figure 5C).

### 3.5 PIR1 N- and C-termini are critical for secretion and cellular localization

Besides their differential, although synergic, involvement in cell wall stability, both PIR-1 and PIR-2 are also structurally different, which imply a potential different way to interact with the cell wall. In addition to the inverted architecture of *N. crassa* PIRs, PIR-1 harbors a Kex2 processing signal, also present in canonical yeast PIRs and associated to the folding and maturation of the protein [18,69,70], and an atypical GPI modification site that, besides the PIR and the Cys-rich domains, could be a third cell wall attachment mechanism [23]. Among yeasts PIRs, a GPI attachment signal has only been reported for *S. cerevisiae* Cwp1p [23]. On the other hand, PIR-2 lacks atypical processing signals, it is devoid of a Cys-rich domain, but contains three PIR domains to bind the wall.

In order to test the functionality of PIR-1 putative processing signals, it was co-translationally fused to *Aequorea victoria* eGFP in three different configurations: N-terminal fusion after the predicted Kex2 processing site (eGFP-PIR-1Kex), C-terminal fusion (PIR-1-eGFP), and to the C-terminus of PIR-1 truncated three amino acids before (T302) the predicted GPI addition site (PIR-1ΔGPI-eGFP) (Figure 6). All *pir-1* and *pir-2 egfp* fusions were targeted to *N. crassa his-3* locus and their expression was controlled by the *Pccg-1* promoter. To visualize the localization of eGFP-tagged PIR-1 and PIR-2, each transformant was directly observed using confocal microscopy.

**Figure 6.**
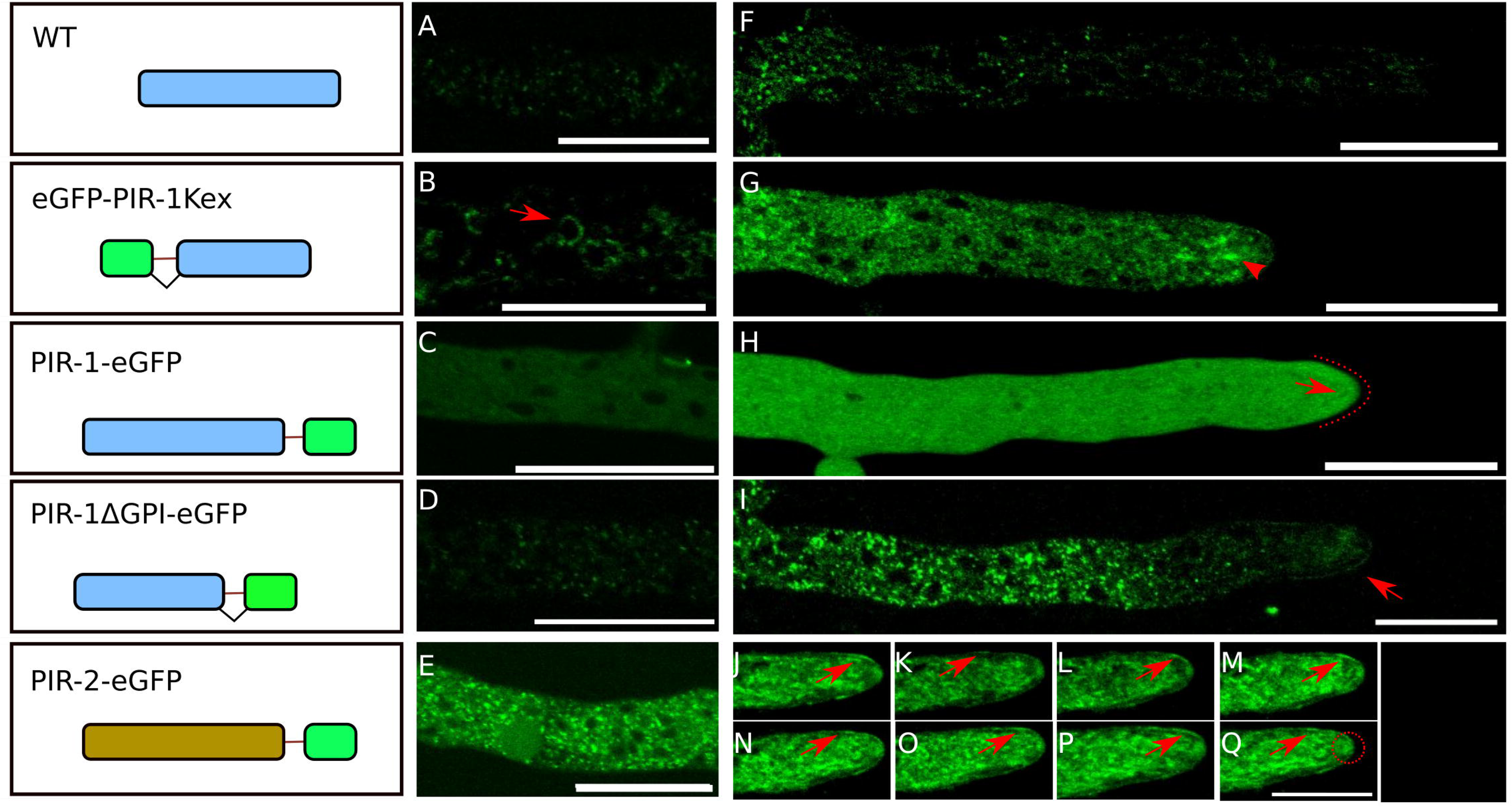
Dissection of *N. crassa* PIRs processing signals. In order to determine the cellular localization and test the functionality of Kex2 processing and GPI addition signals, PIR-1 and PIR-2 were co-translationally fused to eGFP in different positions of their amino acids sequence. As expected, A) distal and F) apical regions of Nc WT (FGSC9013) showed no significant fluorescence. Insertion of eGFP after the predicted Kex2 site of PIR-1 stuck the fusión around nuclei in distal regions of hyphae (B, red arrow), but appeared cytosolic and within small bodies in the apical region (G, red arrow). eGFP C-terminally fused to PIR-1 appeared cytosolic in both distal (C) and apical (H) regions. Nevertheless, a horseshoe-like structure was also observed in the apex (H, red dotted line). eGFP fused to the C-terminus of PIR-1 truncated 3 amino acids before the Ω site appeared in the apical cell periphery as discrete patches (I, red arrow). Finally, eGFP C-terminally fused to PIR-2 (PIR-2-GFP) appeared in small bodies in distal hyphal regions (E), and accumulated in the apical cell periphery (J-Q, red arrows) as well as in the Spitzenkörper (Q, red dotted circle). Scale bar, 20 μm.

#### eGFP-PIR-1Kex

In contrast to Nc *WT* (Figure 6A and F), *N. crassa* expressing eGFP fused to PIR-1 just 3 amino acids after the Kex2 processing site was observed in distal hyphal regions heterogeneously distributed around round structures, putatively nuclei (Figure 6B, red arrow). This signal seemed to cover almost completely those nuclei, but often accumulated in discrete patches over the same structure. Interestingly, in medial and apical regions the perinuclear signal was not observed anymore (Figure 6G); nevertheless, there was an increase of cytosolic fluorescence and the apparition of fluorescent puncta in medial and apical regions (Figure 6G, red arrow).

#### PIR1-eGFP

*N. crassa* expressing PIR1-eGFP showed a strong fluorescence both at the distal and apical cytosol (Figures 6C and H; video S1). Nevertheless, a fluorescent apical horseshoe-like accumulation was observed (Figure 6H, red arrow; video S1), whose fluorescent signal was clearly excluded from the Spitzenkörper (video S1). Considering those images corresponded to a thin optical slide, the 3D projection of the horseshoe-like structure should be a layer of PIR-1-eGFP that covers the cytosolic side of the apical dome.

#### PIR-1ΔGPI-eGFP

In order to further test the functionality of the GPI-addition site, a shortened version of PIR-1 truncated three amino acids before the putative GPI modification site was co-translationally fused to eGFP. This fusion mostly accumulated in small fluorescent bodies in distal and medial regions of the hypha (Figure 6D and I), but also in the apical cell periphery (Figure 6I; video S2). Time series analysis showed a patchy accumulation of PIR-1ΔGPI-eGFP from the apex to more distal regions (up to 20 μm from the apex) of the cell periphery (Video S2). PIR-1ΔGPI-eGFP patches were heterogeneous in size and fluorescence intensity; once they reached the critical distance from the apex the signal vanished.

#### PIR-2-eGFP

C-terminal eGFP-tagged PIR-2 appeared both at distal and proximal regions of the hypha (Figure 6E, J-Q). Unlike PIR-1, PIR-2-eGFP accumulated in the Spitzenkörper (Figure 6J-Q) and also in the cell periphery. There, the signal appeared stronger in discrete regions, although less patchy than PIR-1ΔGPI-eGFP (Figure 6J-Q, red arrows; video S3). As in the case of PIR-1ΔGPI-eGFP, PIR-2-eGFP accumulated in the apical dome up to approximately 20 μm from the apex, behind which the signal disappeared. Times series analysis showed a waiving displacement of the peripheral fluorescent patches from the tip to the medial region of the hypha (Video S3).

## 4 DISCUSSION

After GPI-proteins, PIRs are the second most abundant CWPs [12,85]. Their role is to provide strength and stability to the cell wall either through the formation of covalent bonds between the conserved Q residue of PIR domains and β-1,3-glucans, or the formation of disulfide bonds between the cysteines of the Cys-rich domain and already present CWPs [14,15,22]. Although PIRs have been inferred in filamentous fungi [34], previous phylogenetic, cellular, and biochemical analyses have only been performed on yeast PIRs [25,30]. Here, a comprehensive search of PIR-domains containing proteins in the proteomes of 840 species, representatives of the fungal tree of life, was undertaken to identify the whole diversity of such proteins, but also to better understand their evolutionary history and biological role.

### 4.1 PIRs are an innovation of Ascomycota

PIR-domains containing proteins were found exclusively in Ascomycota and no vestigial PIR domains were identified in other fungal divisions (Figure 1). Previously, Ruiz-Herrera et al. [75] predicted the absence of PIRs in *Ustilago maydis*, a basidiomycete. In a more restricted mass spectroscopy study of *Mucor circinelloides* and *Cryptococcus neoformans* cell wall proteome, PIRs were not detected [86]. In fungi-like *Phytophthora ramorum*, an oomycete, despite its cell wall proteome showed some of the features of ascomycetous CWPs (threonine, serine and proline-rich tandem repeats) also lacked PIRs [87]. PIRs are presumably an innovation of Ascomycota.

Within Ascomycota, some classes were devoid of PIRs, which suggests gain and loss of genes through species evolution. It is also important to notice that within the classes encoding PIRs, the average number of PIRs was maximum among Saccharomycetes species (4.25 proteins/species) followed by Eurotiomycetes (3.4 proteins/species), Dothideomycetes (2.74 proteins/species) and, the lowest, Leotiomycetes and Sordariomycetes, which only contained 1 single protein/species. A biased detection of PIRs in Saccharomycetes because of an imperfect construction of HMMs based in 39 yeast PIRs cannot be ruled out [12]; however, a new HMM built including putative PIR domains from filamentous fungi did not retrieve more sequences (data not shown). On the other hand, although PIRs are bound to the cell wall by an atypical ester linkage that selectively could enrich PIRs in observed ascomycetes [10,14,15], β-1,3-glucan, their bound cell wall polysaccharide, is widely present in fungal cell walls and cannot explain such enrichment. It is well known that structural CWPs evolve faster than enzymatic CWPs so that the fungal cell could adapt to a myriad of environmental challenges [8,11,27]. It seems possible that PIRs, structural CWPs, were a versatile evolutionary solution to changing cell wall destabilizing environments (variable carbon sources, extreme pH, temperature, osmotic pressure) usually experienced by Saccharomycetes, Eurothiomycetes and Dothidiomycetes species [88–91]. This adaptive versatility would be partially provided by a greater number of PIRs in each species (Figure 1), and also by a wide number of PIR domains within such proteins that might provide a differential cell wall stability [16–18,21]. This is particularly relevant for *S. cerevisiae*, regarded as a nomad yeast with no specific niche that, anyway, would need to get adapted to diverse environments [92].

### 4.2 PIRs are a diverse family

PIRs are not an homogeneous group. They grouped in 11 clusters mainly differentiated by the protein architecture (Figure 2). Cluster 1, the most populated, was composed of classical yeast PIRs (cluster 1A) and a new class of filamentous fungi PIRs (cluster 1B; Figure 2B). Both sub-clusters harbor a variable number of PIR domains plus a Cys-rich domain; however, their arrangement is polarized in opposite directions in each one. In cluster 1A, PIR domains are distributed toward the protein N-terminus and the Cys-rich domain is within the C-terminus, while among cluster 1B PIRs, PIR motifs are allocated closer to the C-terminus and the Cys-rich domain is clearly N-terminal (Figure 2C).

The variable number of PIR domains might be the result of an equal number of intragenic PIR domain duplications, which usually occur in the middle of the protein, and mechanistically might be explained by DNA replication slippage [93–97]. Tandem repeat expansion is a mechanism of rapid protein evolution with phenotypic consequences (cell surface variability for fast adaptation to diverse environments) observed in fungi [8,27,98,99], but also in other kingdoms [100]. Inversion of the architecture of filamentous fungi PIRs, with conservation of the protein fold, suggests a circular permutation [101], also a mechanism of protein diversification for better adaptation to changing environments.

Interestingly, the Cys-rich domain (only present in clusters 1A, 1B and 3) contains an embedded and imperfectly conserved PIR-like domain that is the specular image of the *bona fide* PIR domain (Figures 2D and E, S4 and S5). Sequence-reversed protein repetitions have been already reported for immunoglobulins, ferredoxins and membrane proteins [102–105]. Those inverted repetitions are usually arranged around a symmetry axis and are prone to adopt the same fold as homologous direct-sequence domains [106]. Although the evolutionary origin of such inverted repetitions is still obscure, it has been speculated that they arose from a classic gene duplication followed by mutation that led to pseudo-symmetrical inverted repetitions [103,105]. The most conspicuous feature of mirror PIR domains is they are embedded within the Cys-rich domain (Figure S4 and S5). In analogous sequence-reversed domains, a decrease of structural stability regarding the direct domain has been observed, which is mitigated by mutating amino acids that improve the hydrophobic collapse and folding [106]. In a similar way, Cys residues within the mirror PIR domain could stabilize it. Modeling of mirror PIR domain revealed it adopts the same fold as regular PIR domains (Figure S7) and, although empirical confirmation is necessary, it cannot be discarded the former is also functional since key amino acids of the domain (Q, G, D, Q) are strictly conserved (Figure 2A, E, S7). Nevertheless, mirror PIR could also incorporate conformational differences, derived from the reversed sequence, that would provide it with special features that would explain its conservation among class 1A, 1B and 3 PIRs. In fact, the Cys-rich domain of *S. cerevisiae* Pir1p, which also harbors a mirror PIR domain, has been demonstrated as necessary for secretion and correct cell wall localization [107]. Proteins that only contain mirror PIR domains, but not the direct one, could be *bona fide* PIRs, but empirical evidence is still necessary (Table S2). Proteins that only contain a mirror PIR domain also pose the question about which really was the duplicated and inverted PIR domain, the regular PIR domain or the mirror one.

### 4.3 PIRs are disordered proteins, but PIR domains do not necessarily match with that regions

Tandem repeats have been associated with protein’s intrinsically disordered regions (IDR) [62]. Intrinsic disorder buffers the integration or deletion of sequence repeats, which generates evolutionary useful variability without compromising the protein’s core structure [108]. Despite four different predictive platforms, including AlphaFold, confirmed the widespread presence of IDRs in PIRs (Figure 3, S6 and Table S2), PIR domains did not consistently fall within an IDR. Although at least one representative protein of each PIR cluster with PIR domains within an IDR were identified (Figure S6), PIR domains of other conspicuous PIRs (*S. cerevisiae* ScPIR4p, Figure 3) did not match with an IDR. Therefore, PIRs IDRs should have other functions than accommodating tandem repeats. IDRs have also been associated with protein-protein interactions involved in cytoplasmic cell signaling [109]. Beyond their cytoplasmic functions, extracytoplasmic IDRs are able to detect destabilizing defects in the bacterial cell wall which, after signal transduction, helps to keep the wall homeostasis [110] [65]. Widespread presence of IDRs within PIRs, particularly in the opposite terminus to the Cys-rich domain, suggest they could participate in binding modulation to the wall β-1,3-glucan [14,111]. Considering the involvement of PIRs in the response to heat [12], osmotic [112] and antifungals-induced cell wall stress [113], it cannot be ruled out that their IDRs could also play a direct role in the wall structural maintenance. Nevertheless, it has also been proposed that PIR motifs within IDRs could fold into functional two-faced β-helical domains that would allow the formation of covalent bonds with β-1,3-glucans [25].

### 4.4 Despite their inverted architecture, *N. crassa* PIR-1 and PIR-2 are cell wall stabilizers

Except for some *in silico* analysis [34], PIRs had not been studied in filamentous fungi. In yeasts, they have been reported as stabilizers of the cell wall [11,13,17,114,115]□ and mediators of resistance to plant antifungals [113]. Their absence leads to increased cell wall chitin deposition [13,115], increased susceptibility to cell wall synthesis inhibitors [11,17,114], and increased virulence in pathogens that express them, such as *C. albicans* [115]. The inverted architecture of filamentous fungi PIRs, as well as the atypical composition of some of them (Figure 2), raised questions about their role in fungi.

As good representatives of filamentous fungi PIRs, single knockout Nc Δ*pir-1* and Nc Δ*pir-2* mutant strains were analyzed to infer their biological role (Figure 3). They did not show any deleterious phenotype and revealed single mutations do not compromise neither cell viability nor growth rate (Figure 4). Because a synergic effect of the double Nc Δ*pir-1/*Δ*pir-2* mutant could not be ruled out, the double knockout mutant was constructed. However, a Nc Δ*pir-1/*Δ*pir-2* homokaryotic strain could not be isolated, probably because of the joint importance of both PIR-1 and PIR-2 proteins in *N. crassa* (Fig S2). As expected, the heterokaryotic Nc Δ*pir-1/*Δ*pir-2* strain showed a synergic effect on hyphal growth rate (Figure 4), which, together with the features of PIRs, indicated a cell wall defect. A similar behavior has been observed in *S. cerevisiae* and *C. albicans* single PIR mutants, which did not show a clear phenotype [18,30]; however, the double *S. cerevisiae* Δ*pir1*/Δ*pir2* mutant exhibited a slow growing phenotype [12]. Moreover, simultaneous deletion of PIR1 and PIR32 in *C. albicans* led to a hyper filamentous growth [114–116].

To further explore the physiology of *N. crassa* PIRs, Nc Δ*pir-1*, Nc Δ*pir-2*, and Nc Δ*pir-1*/Δ*pir-2* mutant strains were subjected to stress assays with CFW, NaCl, and heat (Figure 5), stressors that exacerbate cell wall damage. As observed with growth rate, CFW (chitin synthesis inhibitor) and NaCl (osmotic stressor) effect was not different between Nc WT, Nc Δ*pir-1*, and Nc Δ*pir-2*; nevertheless, the double mutant Nc Δ*pir-1*/Δ*pir-2* showed around 50% growth inhibition versus 20% of single mutants in presence of 1 mg/ml CFW, and 75% growth inhibition versus around 50% of single mutants in 4% NaCl. This indicated that the cell wall damage inflicted by any of both PIRs is not enough to compromise the wall stability; however, the double mutant did. In *S. cerevisiae*, single mutants of PIR-1p, PIR-2p, PIR-3p and PIR-4p led to increased sensitivity to CFW and Congo red (another cell wall synthesis inhibitor), and the effect of those stressors was stronger when multiple PIR mutants were produced [11].

Unlike Nc Δ*pir-1*, response of Nc Δ*pir-2* to heat stress suggests it could be part of the cellular heat shock response. In *S. cerevisiae*, Pir2p/HSP150 is upregulated in response to heat [85,117]. Noteworthy, *C. albicans* PIR1 and PIR32 mutants did not show any heat shock related phenotype [30,115]. This indicates that PIRs are not functionally redundant, as previously stated.

Taking these data together, it can be suggested that the main role of PIRs in cell wall maintenance is conserved in both yeast and filamentous fungi, despite the inverted protein architecture of the latter and the functional differences that PIRs present.

### 4.5 Signals processing and localization of *N. crassa* PIR-1 and PIR-2

The atypical architecture and presence of multiple processing signals of PIR-1, belonging to cluster 1B PIRs, posed questions about their maturation, secretion and cell wall binding. Besides its PIR and Cys-rich domains, PIR-1 also contains a Kex2 processing site and a GPI addition signal (Figure 6, Table S2). GPI-containing PIRs have been previously reported in *S. cerevisiae* [23] and *C. glabrata* [33]. Unlike yeast GPI-PIR proteins, filamentous fungi PIRs additionally have a Cys-rich domain.

eGFP-tagging of PIR-1 just after the Kex2 processing site led to an accumulation of PIR-1 in circular structures that resembled the perinuclear endoplasmic reticulum (ER) (Figure 6b) [118,119]. Processing of Kex2 site in a number of yeast proteins (including PIRs) at the trans Golgi network is necessary for correct maturation and folding [15,69,114,120–122]. Fluorescence allocation of eGFP-PIR-1Kex, in contrast to C-terminal eGFP fusions to PIR-1 (Figure 6), suggests eGFP-PIR-1Kex secretion is impaired at a step before Kex2 processing, which lead to an increase of the fusion residence time in the ER, allowing its detection (Figure 6B). However, an analogous fusion of *S. cerevisiae* Pir2p/HSP150 with mRFP did not show such mislocalization [107], which suggests the inverted architecture of filamentous fungi PIR proteins could led to an increased structural sensitivity of *N. crassa* PIR-1 to N-terminal GFP-tagging. Besides the Kex2 site, PIR-1 also harbors a GPI-addition signal that is particularly interesting since it could be a third cell wall attachment mechanism in addition to cell wall bonding via PIR and Cys-rich domains. To assess the functionality of the GPI-addition site, a C-terminal fusion of PIR-1 with eGFP was constructed (Figure 6C and H). *N. crassa* PIR-1-eGFP expressing strain showed cytoplasmic fluorescence (Figure 6C and H), which suggests the GPI signal is functional and the GFP-tagged C-terminal signal peptide is released to the cytosol. Despite GPI-proteins C-terminal signal peptide harbors a hydrophobic domain that anchor it to the ER membrane, it has been demonstrated that fused exogenous peptides can influence the position and stability of the C-terminal signal peptide [123], thus it is possible that GFP-tagged C-terminal signal peptide might be unstable enough that it is ultimately released to the cytosol.

Interestingly, PIR-1 truncated three amino acids before the Ω-site and fused to eGFP, PIR-1ΔGPI-eGFP (Figure 6I, video S2), as well as PIR-2 C-terminally fused to eGFP (Figure 6J-Q), accumulated as discrete patches in the apical cell periphery from the very tip up to approximately 20 μm backwards (Videos S2 and S3). A similar patchy allocation pattern has been reported for *S. cerevisiae* Pir1p-eGFP and Pir2p (Hsp-150)-myScarlet [25,107], and also for *S. cerevisiae* Pir1p, Pir2p, and Pir4p detected by immunofluorescence [124–126]. *S. cerevisiae* Pir4p mainly appeared in the cell surface of growing buds, which suggested it could be associated with newly synthesized cell wall [124], while Pir1p-eGFP was observed in bud scars [107]. Although additional experiments are necessary to assess the relationship of PIR-1 and PIR-2 localization around the hyphal apex with cell polarity, disruption of cell polarity and bud selection markers (BNI1p, SPA2p, PEA2p, AXL2p, BUD3p, BUD8p, RGA1p, BEM1p, and BEM2p) in *S. cerevisiae* did not affect the localization of Pir1p, indicating Pir1p localization is not associated to cell polarity or bud scar selection [107]. The irregular distribution of PIRs on the cell surface of both yeast and filamentous fungi suggests they could be associated with discrete cell wall domains from which they exert their stabilizing effect.

All collected observations indicate PIRs are an innovation of Ascomycota and their presence coincides with those fungal classes that experience changing environments that challenge their cell walls stability. Despite the inverted architecture of filamentous fungi PIRs, they keep the same fold and cell wall stabilizing role of classical yeast PIRs. In addition to PIR and Cys-rich domains, a potentially active mirror PIR domain (present in both yeast and filamentous fungi PIRs) and a GPI-modifying signal could be alternative mechanisms to anchor PIRs to the wall. It is intriguing the role of widespread intrinsic disorder among PIRs; however, the accumulation of PIRs in discrete patches or plugs suggest IDRs undergo disorder-order transitions that might mediate PIRs interaction (Figure 7). AlphaFold multimerization modeling predicts both PIR-1 and PIR-2 interact with themselves via their Cys-rich domains, while PIR domains fold and expose the glutamine residue that potentially interacts with β-1,3-glucans (Figure 7). It is also intriguing how PIRs manage to stabilize the cell wall without forming a more intuitive homogeneous network that covers it. Instead, they accumulate in plugs, whose composition (either a single PIR or a mixture of both) and mode of action will be the subjects of an upcoming investigation (Figure 7).

**Figure 7.**
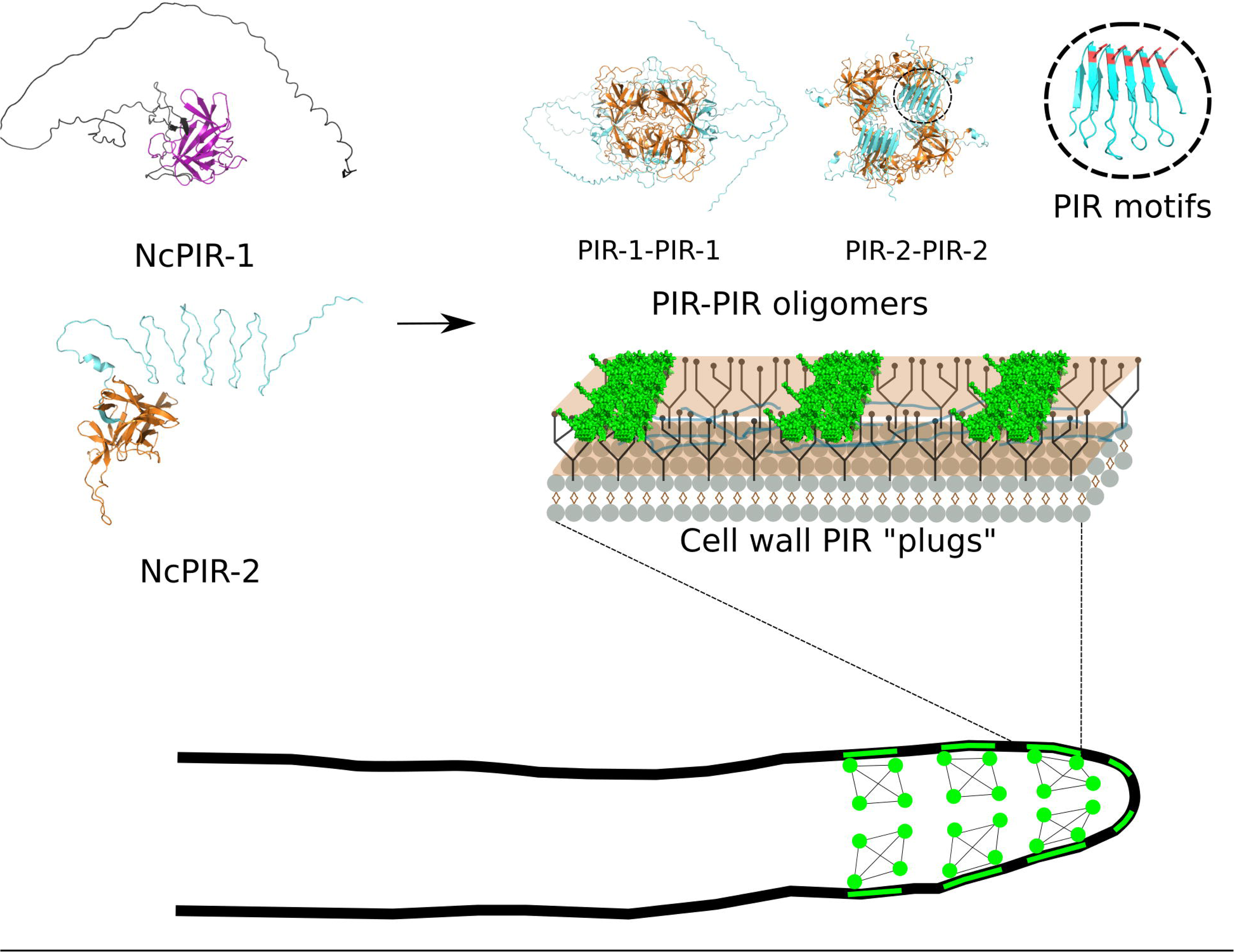
The role of PIR-1 and PIR-2 in the cell wall of *N. crassa*. *N. crassa* PIR-1 and PIR-2, representatives of filamentous fungi PIR proteins (cluster 1B), are cell wall stabilizers as classical yeast PIR proteins (cluster 1A), despite their inverted architecture. PIR-1 and PIR-2 accumulate as “plugs” in the apical cell periphery. Multimerization predictions suggest those plugs could be PIR-1 and/or PIR-2 protein complexes formed after disorder-order transitions of their intrinsically disordered regions. How those “plugs” provide stability to the cell wall will be the subject of future research.

## 6 AUTHOR CONTRIBUTIONS

PM-S performed experimentation. PM-S and JV designed experiments, analyzed results, and drafted and edited the manuscript. OAC-N provided confocal microscopy support. AP-S made helpful suggestions on bioinformatic analysis. JV conceptualized research and acquired financial support. All authors critically reviewed and commented on the manuscript, and approved the final version.

## 7 ACKNOWLEDGMENTS

We thank Meritxell Riquelme and Diego Delgado for kindly hosting Paul Montaño-Silva during his stay at the Department of Microbiology and LNMA, CICESE, in Ensenada. We also appreciate the lab management work of Flor García Niño and the support of CIATEJ administrative staff.

This work was supported by grants from SENER-CONACYT Mexico (245750), Ciencia de Frontera, CONAHCYT-Mexico (2019-552259), and COECyTJAL-FODECIJAL (8186-2019). PM-S (CVU 730994) was recipient of MSc and PhD fellowships from CONAHCYT-Mexico.

