## Supplemental Materials for "Cell wall-resident PIR proteins show an inverted architecture in *Neurospora crassa*, but keep their role as wall stabilizers"

RUNNING TITLE: A new class of fungal PIR proteins

Paul Montaño-Silva^1^, Olga A. Callejas Negrete^2^, Alejandro Pereira-Santana^3^ and Jorge Verdín^1,^*

^1^Biotecnología Industrial, CIATEJ-Centro de Investigación y Asistencia en Tecnología y Diseño del Estado de Jalisco, Zapopan, JAL, Mexico.

^2^Departamento de Microbiología, CICESE-Centro de Investigación Científica y de Educación Superior de Ensenada, Ensenada, BC, Mexico.

^3^CONAHCYT-Centro de Investigación y Asistencia en Tecnología y Diseño del Estado de Jalisco, Sede Sureste, Merida, YUC, Mexico.

**Supplementary Material**

**Materials and methods**

*Molecular construction of N. crassa his-3 targeted assemblies.* Unless a different template is indicated, DNA fragments were PCR amplified from *N. crassa* FGSC 9013 genomic DNA. *egfp::v5* and *v5::egfp* were amplified from pre-assembled plasmids pGEM::NCW3 (*ncw-3::V5::gfp::5'his-3*) and pGEM::ACW1 (*gfp::V5::acw-1::5'his-3*), respectively, kindly donated by Ana Sofía Ramírez-Pelayo [127]. To C-terminally fuse eGFP to PIR-1, *Pccg-1::pir-1::v5::gfp* (PIR-1-eGFP) and *Pccg-1::pir-1_1-1027_::v5::egfp* (PIR-1ΔGPI-eGFP) the respective constructs were built (Figure S1 A, B, E). For *Pccg-1::pir-1::v5::egfp, pir-1* was PCR amplified with CPIR 1-4 and CPIR 1-5 primers, while *v5::egfp* was amplified using CPIR 1-6 and r_GFP-BstBI primers. For C-terminal eGFP fusion to PIR-1 without GPI signal, *pir-1*_1-1027_ was PCR amplified using CPIR 1-4 and rPIR1-GPI primers, and *v5::egfp* was amplified using f_v5:GFP_GPI and r_GFP-BstBI primers. N-terminal fusion *Pccg-1::pir1_1-244_::egfp::v5::pir-1_245-1155_* (eGFP-PIR-1Kex) was constructed as follows: *pir-1*_1-244_ and *pir-1_244-1155_* were amplified using CPIR 1-4 and R_PIR 1Kex, and FPIR1s/PS+Kex and PIR1-BstBI primers, respectively. *egfp::v5* was amplified with FGFP-Kex and rGFP:V5Kex primers (Figure S1 C, E). For C-terminal fusion of PIR-2 with eGFP, *pccg-1::pir-2::v5::egfp*, *pir-2* was amplified using XbaI-PIR2 and CPIR 2-3 primers, while *v5::egfp* was amplified with CPIR 2-4 and r_GFP-BstBI (Figure S1 D, E).

To construct each assembly, the corresponding fragments were mixed together in a one-step assembly reaction using the NEBuilder Assembly Kit (New England Biolabs) as follows: 0.1 pmols each fragment was added to the reaction in a final volume of 10 uL and incubated for two hours at 50 °C. Then, each product was PCR amplified using the corresponding primers: XbaI-PIR-1 and PIR1-BstBI for eGFP-PIR-1Kex; XbaI-PIR1 and r_GFP-BstBI for C-terminal fusions of PIR-1, and XbaI-PIR2 and r_GFP-BstBI for PIR-2 assembly. In all cases, XbaI and BstBI restriction sites were added to the 5’ and 3’ ends, respectively. Because of the low PCR efficiency, all four cassettes were first ligated into pGEM-T Easy Vector (Promega) and cloned in *E. coli* TOP10 (ThermoFisher) getting the plasmids pGEM::NPIR1Kex, pGEM::CPIR1, pGEM::CPIR1ΔGPI and pGEM::CPIR2 (Table 1). All plasmids were purified by miniprep (GeneElute Miniprep Kit, Sigma) and sequenced. After that, the four resulting plasmids were double digested with XbaI and BstBI restriction enzymes (Anza, ThermoFisher) to release the expression cassettes *pir-1::v5::gfp, pir-1_1-1027_::v5::egfp, pir1_1-244_::egfp::v5::pir-1_245-1155_,* and *pir-2::v5::egfp*, which were subsequently ligated into pMF272 using T4 DNA ligase (Promega) and cloned in *E. coli* TOP10 getting pPMS04, pPMS05, pPMS07 and pPMS09 plasmids, respectively (Figure S1 E).

*Molecular characterization of N. crassa knockouts. N. crassa* FGSC 16451 (*Δpir-1*) and FGSC 17985 (*Δpir-2)* were molecularly confirmed by two genomic PCR reactions. The first was targeted to amplify either *pir-1* (primers FPIR1 and RPIR1) or *pir-2* (FPIR2 and RPIR2) to demonstrate their absence in KO strains (Figure S2 A). The second reaction amplified the *hph* gene that replaced *pir-1* or *pir-2* using primers that hybridizes to *hph* (Fhph) and the 3’ flank of either *pir-1* (RP1-h) or *pir-2* (RP2-h) (Figure S2 B). The double KO strain *N. crassa* *Δpir-1Δpir-2* (NcPMS01) was confirmed by genomic PCR with two primers that hybridize to *natR* gene and *3’pir-2 flank* (2PIR 2-3 and 2PIR 2-4) which amplified the insertion (915 bp) of the assembly (*5’pir-2 flank::natR::PtrpC::3’pir-2 flank)* into the *N. crassa* FGSC 16451 (Δ*pir-1*) genome (Figure S2 C).

*Molecular characterization of N. crassa his-3 targeted transformants. pir-1::v5::gfp, pir-1_1-1027_::v5::egfp, pir1_1-244_::egfp::v5::pir-1_245-1155_* and *pir-2::v5::egfp* were directed to the *N. crassa* *his-3* locus using pMF272 vector. All cassettes were expressed under the control of the *Pccg-1* promoter. Two primers that hybridize the *his-3* locus and its 3’ flank (HisFlakF and UniBstBIR) were used to molecularly confirm the insertion of the assemblies into *N. crassa* genome (Figure S3 A). Because the strains were heterokaryon, two PCR bands were expected: one from the WT untransformed nuclei (1461 bp) (Figure S3 B and C, lanes 2-6), and the other from the transformed ones (around 3,000 bp; Figure S3 B and C, lanes 3-6).

**TABLES**

**Table S1**. Primers used in this work.

**Table S2**. Clustering of fungal PIRs.

**FIGURE LEGENDS**

**Figure S1.** **Assemblies construction for *his-3* targeted transformation of *N. crassa*.** All assemblies (A, B, C, D) were constructed in three general steps. Step one was focused on amplifying each DNA fragment of the assemblies. In step two, each PCR amplicon was mixed together for a one step Gibson assembly reaction using NEBuilder (*New England Biolabs*). Finally, in step three, assembled fragments were PCR amplified to add 5’ XbaI and 3’ BstBI restriction sites to finally clone each cassette into pMF272 (E).

**Figure S2. Molecular confirmation of *N. crassa* KO strains *Δpir-1*, *Δpir-2*, and *Δpir-1/Δpir-2*.** A) Lanes 1, 3 and 5, PCR amplification of *pir-1* from *N. crassa* WT (FGSC 9013), *Δpir-1* and *Δpir-2* strains gDNA using FPIR1 and RPIR1 primers. As expected, *pir-1* did not amplify from *N. crassa* *Δpir-1*. Lanes 2, 4 and 6, PCR amplification of *pir-2* from *N. crassa* WT, *Δpir-1* and *Δpir-2* strains gDNA using FPIR2 and RPIR2 primers. As expected, *pir-2* did not amplify from *N. crassa* *Δpir-2.* B) Lane 1 and 2, PCR amplification of *hph* used to knock out *pir-1* and *pir-2* from *N. crassa* *Δpir-1* and *Δpir-2* gDNA with a pair of primers that hybridized in *hph* (Fhph) and the 3’ flank of either *pir-1* (RP1-h) or *pir-2* (RP2-h), respectively. As expected, two amplicons of 2031 bp and 2039 bp, respectively, were obtained. C) Lane 1, PCR amplification of *PtrpC::natR* (915 bp) from NcPMS01 strain gDNA using 2 PIR 2-3 and 2 PIR 2-4 primers. Lane 2, *N. crassa* WT gDNA was used as negative control and, as expected, *PtrpC::natR* did not amplify.

**Figure S3.** **Molecular confirmation of *his-3* targeted *N. crassa* strains.** A) *his-3* targeted transformants were molecularly confirmed by genomic PCR targeting two primers to the *his-3* locus (HisflankF and UniBstBIR primers). As negative control no gDNA was added to the reaction (lane 1). FGSC 9717 gDNA strain was used as positive control (lane 2). Lanes 3-6, show PCR amplicons that confirmed the insertion of the expression cassette for each heterokaryotic *N. crassa* strain: lane 3, NcPMS04 (2977 bp); lane 4, NcPMS05 (2977 bp); lane 5, NcPMS07 (2905 bp); and lane 6, NcPMS09 (3044 bp), respectively.

**Figure S4. Multiple sequence alignment of C- and N-terminal Cys-rich domains of a representative selection of PIRs from clusters 1A and 1B, respectively.** C-terminal Cys-rich domain of yeasts PIRs is the canonical one and it is composed of four Cys residues (C_1_-x(64)-C_2_-x(16)-C_3_-x(12)-C_4_), all of them located after the Kex2 processing site. On the other hand, the N-terminal Cys-rich domain of filamentous fungi cluster 1B PIRs contains two additional Cys residues, one located before the Kex2 processing site (C_-1_), which is not conserved in the mature protein, and a second additional Cys residue (C_x_) located between the canonical C_1_ and C_2_. A discrete sequences group within cluster 1B does not contain C_1_ and C_4_.

**Figure S5.** **PIRs cluster 3 is expanded with an amino acids insertion of variable length.** Multiple sequence alignment of PIRs cluster 3 revealed an amino acids insertion of up to 47 residues between Cys-1 and Cys-2 of the Cys-rich domain. The black dotted square highlights the single, N-terminal, PIR domain of cluster 3, while the red dotted square notices the amino acids insertion between Cys-1 and Cys-2 of the Cys-rich domain characteristic of this cluster.

**Figure S6.** **Intrinsic disorder prediction in PIRs**. Intrinsic disorder was inferred from three different predictors, IUPRED3 (blue line), ESpritz (red line), FIDpnn (green line) for a representative PIR from each cluster (C1a, C1b, C2, C3, C4, C5 and C6). In addition, PIR domain allocation for each protein is shown (lower line, turquoise blocks). Only in some cases (C1a, C1b, C5 and C6), PIR domain colocalized with intrinsic disorder inferred by at least two predictors.

**Figure S7. AlphaFold prediction of *N. crassa* PIR-1 structure.** A) Tertiary structure prediction of Nc PIR-1 highlighting the direct PIR domain (green arrows) and the mirror-PIR (brown arrows) within it. Besides the structured core (brown surface), most of Nc PIR-1 is intrinsically disordered (light green). B) PIR domain structures as a β-hairpin; amino acids composition, their hydrophobicity and propensity to form a β-turn (GDG). C) Model of the mirror PIR domain structuring as a β-hairpin and amino acids composition, their hydrophobicity and propensity to form a β-turn. The amino acid segment comprising the mirror PIR domain was parallelly inverted in order to align it with the sequence of the direct PIR domain. Asterisks indicate conserved glutamine residue (colored pink) that is used to anchor PIRs to β-1,3 glucans. For β-turn propensity, amino acids in red denote the highest propensity to form a loop, while the blue ones denote the least. The most hydrophobic amino acids are coloured in red, while the most hydrophilic ones are coloured in blue. Underlined amino acids are related to loop formation within the β-hairpin structure.

**Video S1. Live imaging of *N. crassa* NcPMS04 (*Pccg-1::pir-1::v5::egfp*).** PIR-1-eGFP was expressed under the control of the *Pccg-1* promoter. PIR-1-eGFP accumulated mainly in the cytosol. A horseshoe-like accumulation at the apical dome, excluded from the Spitzenkörper, was observed. This video was edited with FIJI. Scale bar, 10 μm.

**Video S2. Live imaging of *N. crassa* NcPMS05 (*Pccg-1::pir-1_1-1027_::v5::egfp)*.** PIR-1ΔGPI-eGFP was mostly accumulated in small fluorescent bodies in distal and medial regions of hyphae. Fluorescent patches were observed in the apical cell periphery. The video was edited with FIJI. Scale bar, 10 μm.

**Video S3.** **Live imaging of *N. crassa* NcPMS09 (*Pccg-1::pir-2::v5::egfp).*** PIR-2-eGFP accumulated in the Spitzenkörper and in the cell periphery. Fluorescent, waving, patches were observed in the apical dome up to approximately 20 μm from the apex. Scale bar, 10 μm.
