## Supplemental Table S1 for "Cell wall-resident PIR proteins show an inverted architecture in *Neurospora crassa*, but keep their role as wall stabilizers"

***Table S1. Primers used in this work***

| **Primers** | **Sequence (5’ to 3’)** | **General Tm (°C)** |
| --- | --- | --- |
| **2 PIR 2-1** | ATCGATCGACTTGCACACGAGACTGTC | 64 |
| **2PIR 2-2** | CCAAAAAATGCTGAAAATCTGTGGATGGGATGG | 64 |
| **2PIR-2-3** | CACAGATTTTCAGCATTTTTTGGGCTTGGC | 65 |
| **2PIR 2-4** | CTCTTCCTTTCAGGGGCAGGGCATG | 65 |
| **2 PIR 2-5** | GCCCCTGAAAGGAAGAGCCCAAGCC | 67 |
| **2 PIR 2-6** | ATCGCTCATGAGAGGAAAGCAAGCAGC | 65 |
| **XbaI-PIR-1** | AGGTCTAGAATGAAGCAATATCAGATCCTGACTCTTC | 63 |
| **XbaI-PIR-2** | AGGTCTAGAATGAAGCAATATCAGATCCTGACTCTTC | 63 |
| **CPIR 1-4** | AACCGTCAAAATGAAGCAATATCAGATCCTGACTC | 65 |
| **CPIR 1-5** | GGTTTGCCGATGAACCGCATGCCAC | 67 |
| **CPIR 1-6** | GGTTCATCGGCAAACCGATTCCGAAC | 64 |
| **CPIR 2-3** | GTTTGCCGAACTGGCCATCGTTGC | 65 |
| **r_GFP-BstBI** | AGCTCGTTCGAATTACTTGTACAGCTCGTC | 64 |
| **rPIR1-GPI** | GGTTTGCCGGTGATCACGACGCTGG | 68 |
| **f_V5:GFP_GPI** | GTGATCACCGGCAAACCGATTCCGAACC | 67 |
| **R_PIR 1Kex** | CCTTGCTCACGACCTGCTGTTGTACGG | 67 |
| **FGFP-Kex** | GCAGGTCGTGAGCAAGGGCGAG | 66 |
| **rGFP:V5Kex** | TGGCACTGGCGGGTGCTATCCAG | 67 |
| **FPIR1s/PS+Kex** | CACCCGCCAGTGCCATGGC | 66 |
| **PIR1-BstBI** | AGCTCGTTCGAATCAGATGAACCGCATG | 65 |
| **CPIR 2-4** | GCCAGTTCGGCAAACCGATTCCGAAC | 66 |
| **HisFlankF** | TTGATTGACAGCGAACGAAACC | 58 |
| **UniBstBIR** | TCAGCATCCGTCTTGAGC | 54 |
| **FPIR1** | GATCCTGACTCTTCTTGCCTGC | 59 |
| **RPIR1** | AGATGAACCGCATGCCACC | 59 |
| **FPIR2** | TCTTTCACCTTGTCCGGCG | 58 |
| **RPIR2** | GAACTGGCCATCGTTGCTTTG | 59 |
| **Fhph** | CACTGACGGTGTCGTCCATC | 59 |
| **RP1-h** | CACGCCCCACTTCTTCGG | 59 |
| **RP2-h** | CCGATCAGGAGACGAAGGCTG | 59 |
| **1PIR 1-2** | GCTGTACAAGTAAGTCGACGTTAACTGGTTCCC | 65 |

*Signal peptides synthesized as oligomers.
