## Supplementary figures and images for "Cell wall-resident PIR proteins show an inverted architecture in *Neurospora crassa*, but keep their role as wall stabilizers"

### Supplemental Figure S1

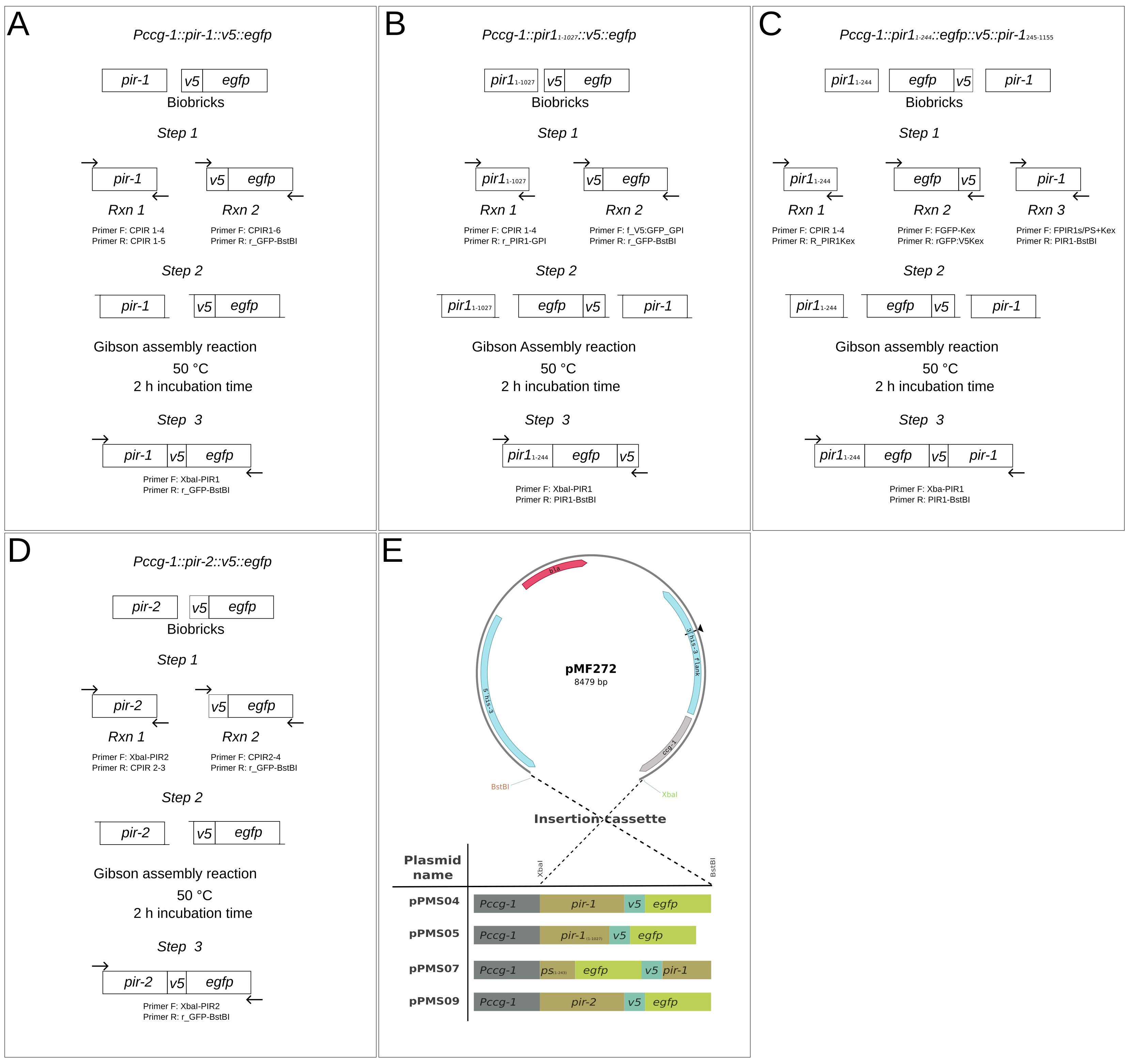

### Supplemental Figure S2

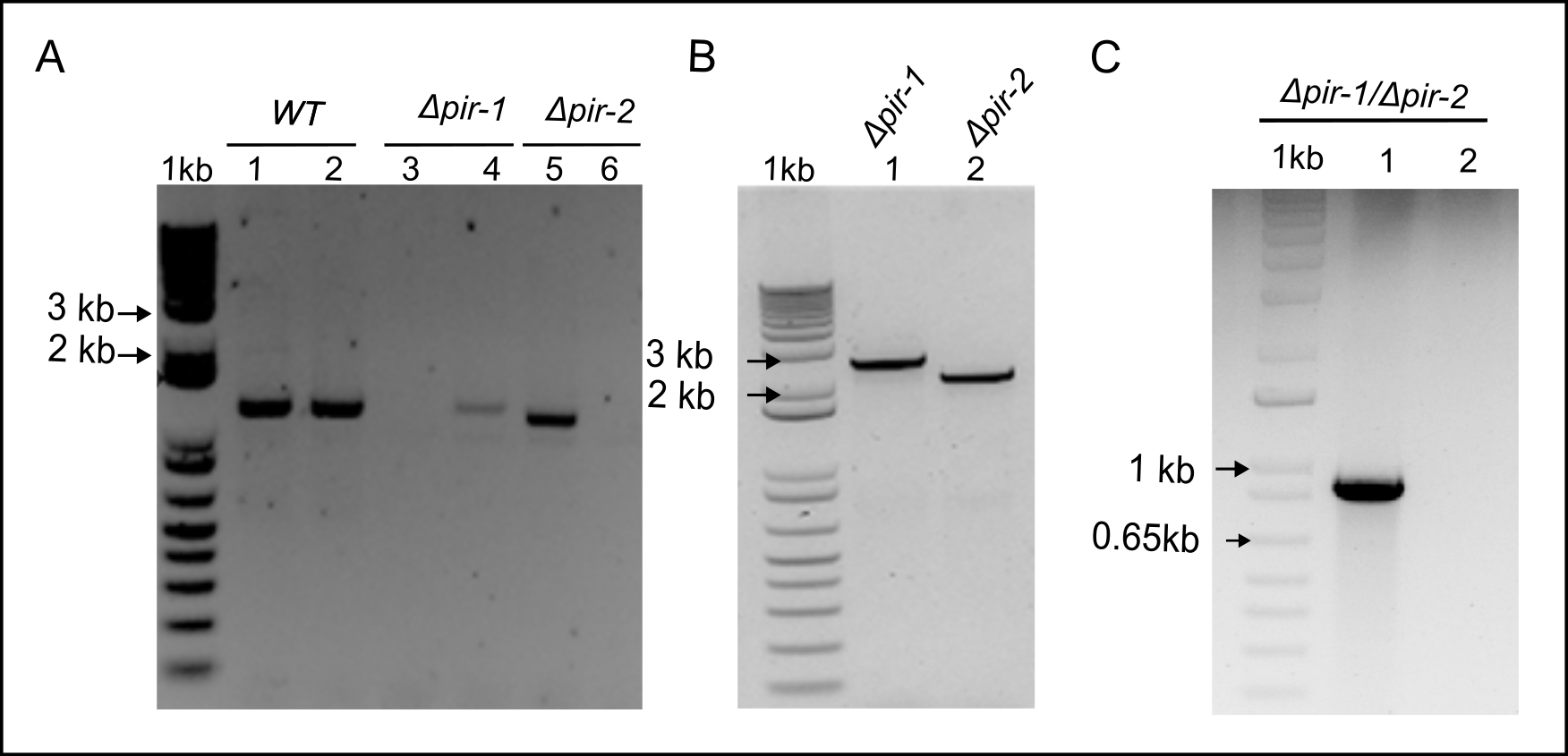

### Supplemental Figure S3

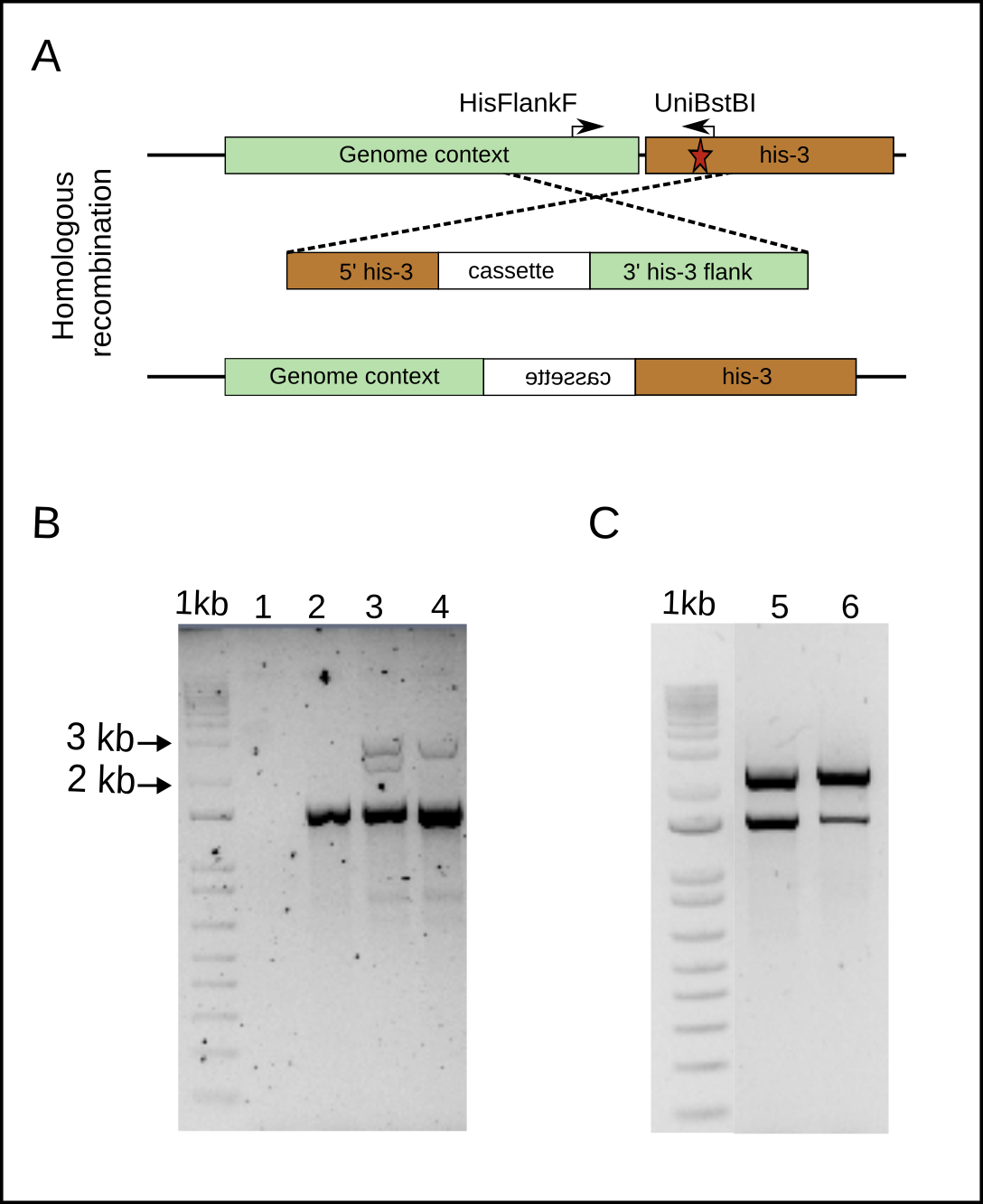

### Supplemental Figure S4

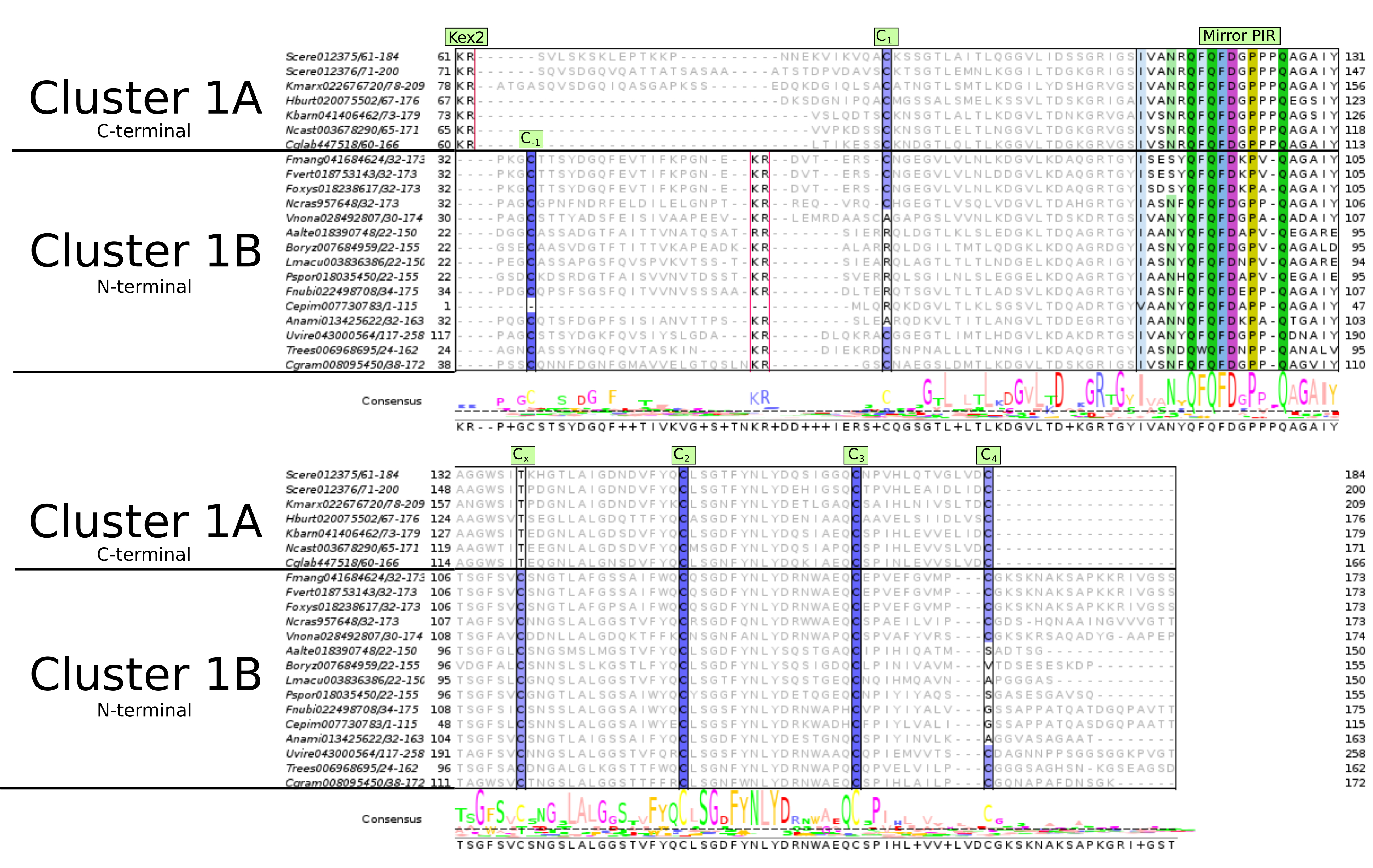

### Supplemental Figure S5

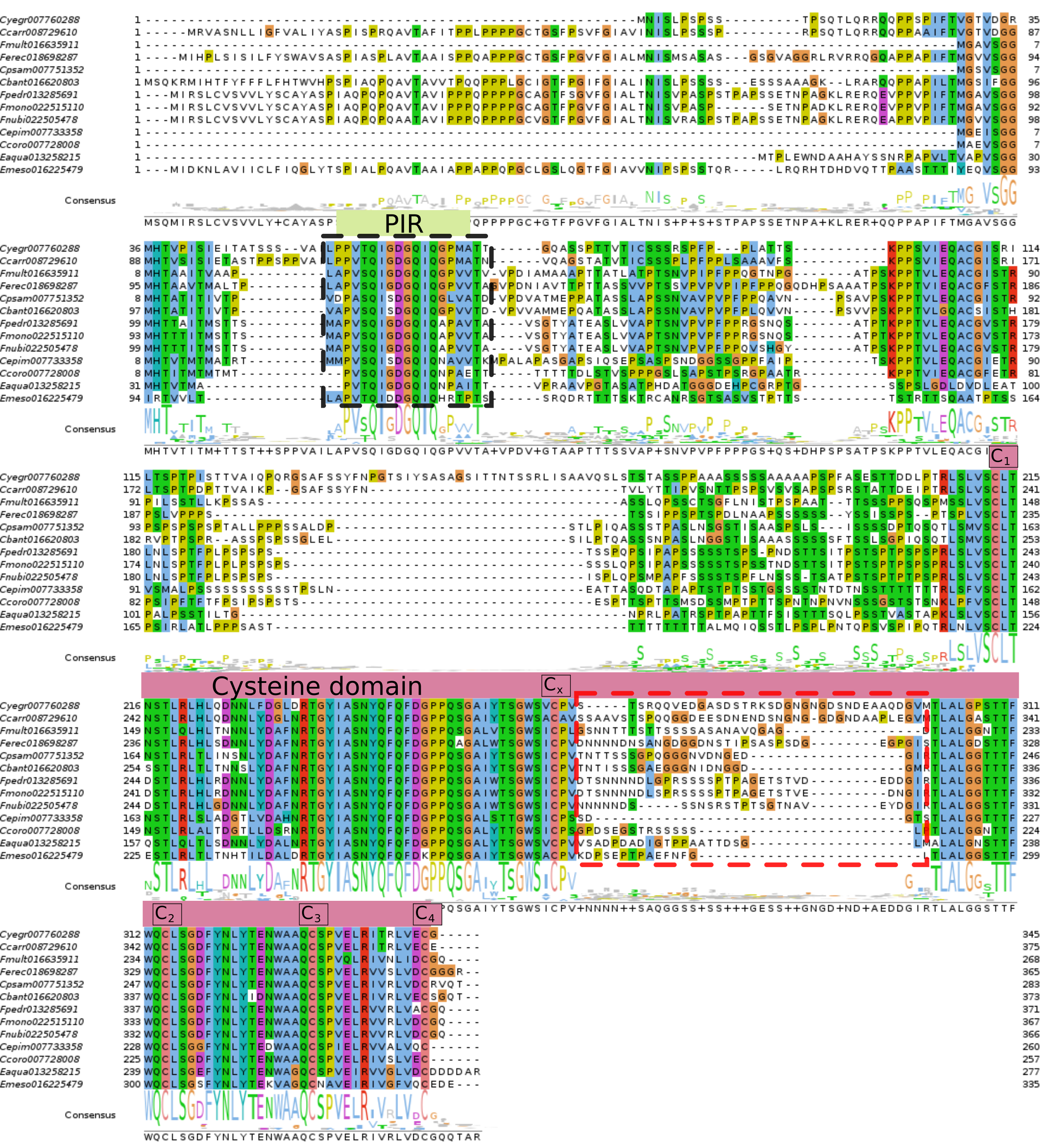

### Supplemental Figure S6

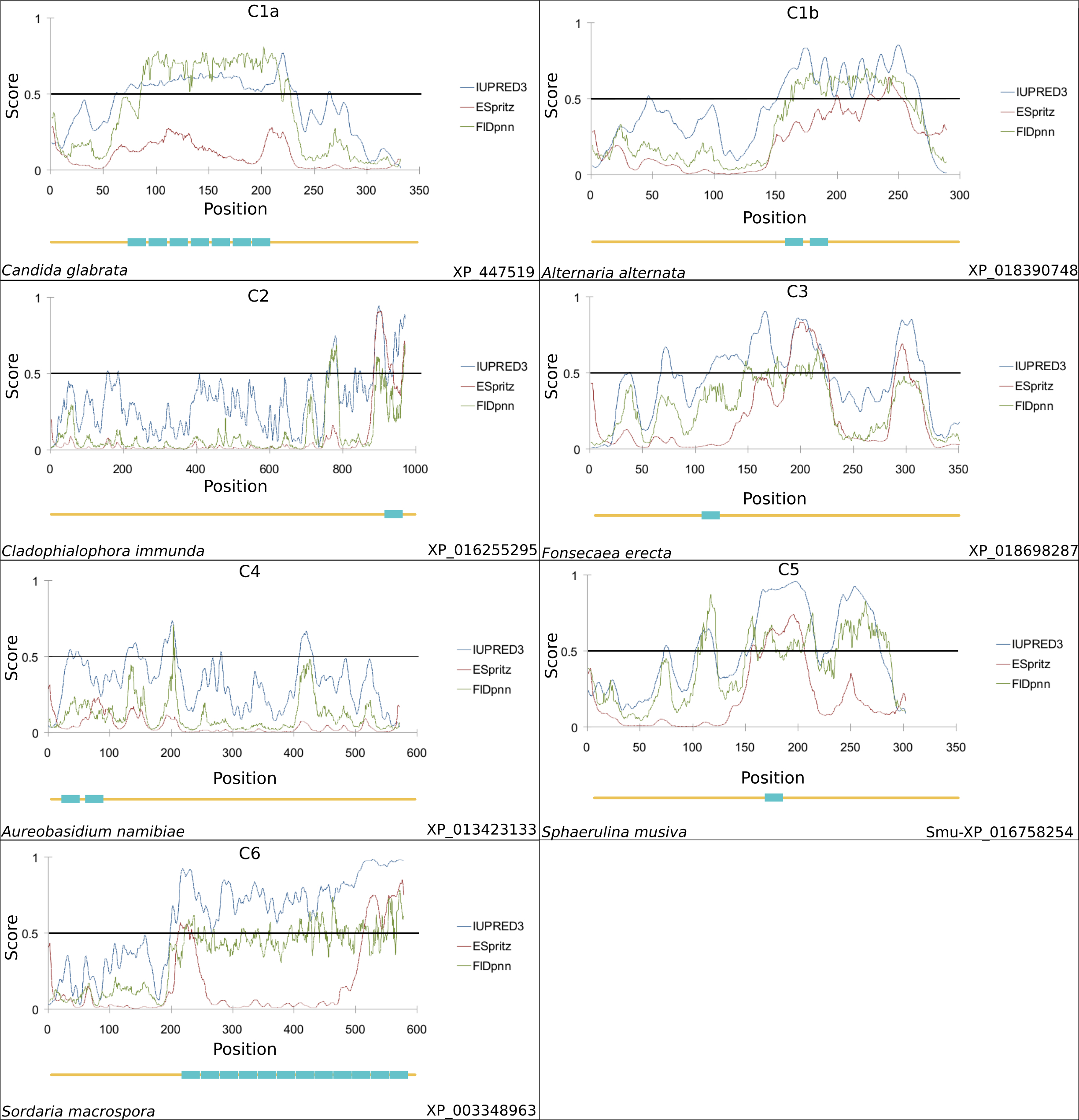

### Supplemental Figure S7

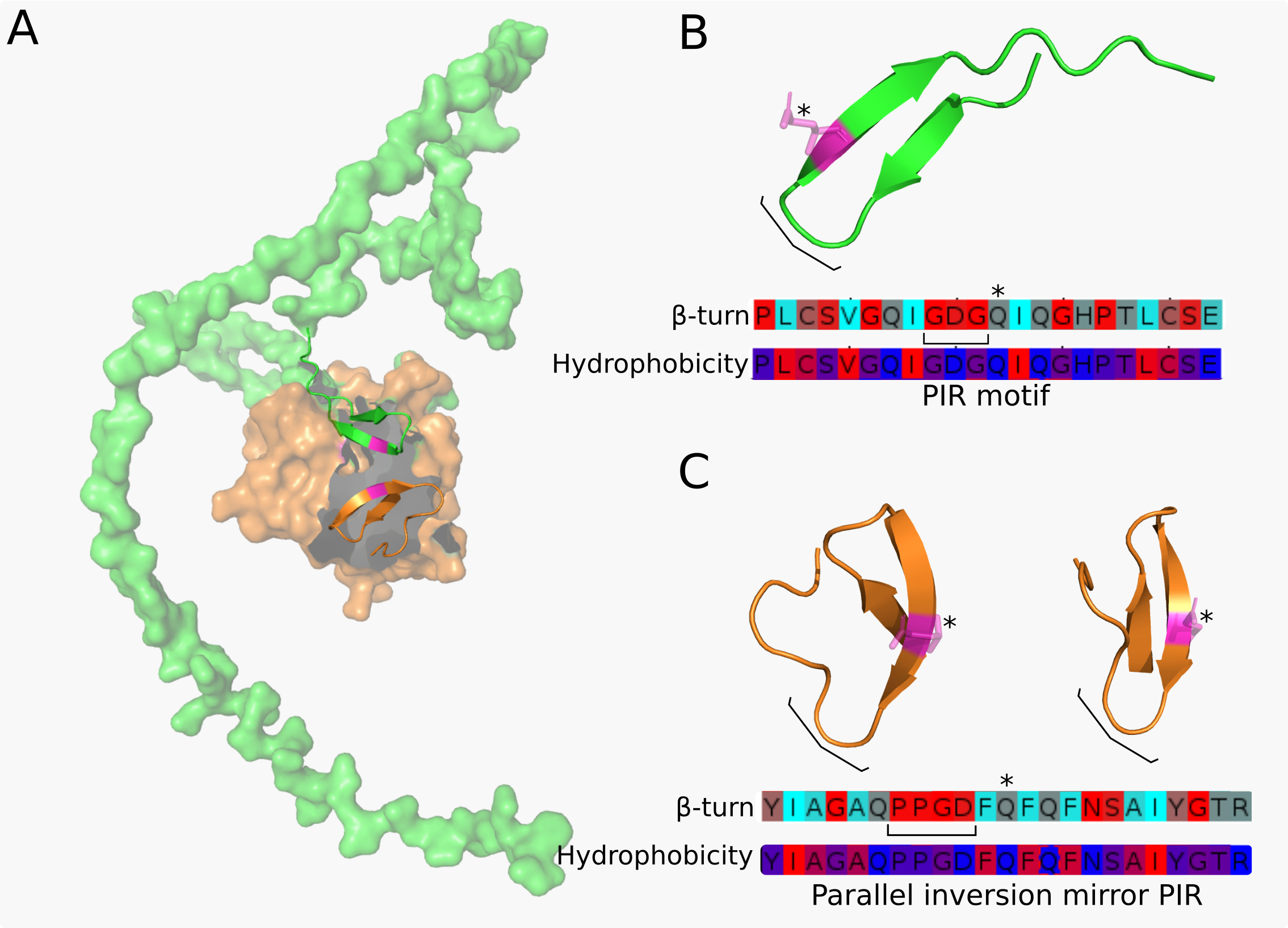
